# Autophagy suppression in DNA damaged cells occurs through a newly identified p53-proteasome-LC3 axis

**DOI:** 10.1101/2024.05.21.595139

**Authors:** D’Feau J. Lieu, Molly K. Crowder, Jordan R. Kryza, Batcha Tamilselvam, Paul J. Kaminski, Ik-Jung Kim, Yingxing Li, Eunji Jeong, Michidmaa Enkhbaatar, Henry Chen, Sophia B. Son, Hanlin Mok, Kenneth A. Bradley, Heidi Phillips, Steven R. Blanke

## Abstract

Macroautophagy is thought to have a critical role in shaping and refining cellular proteostasis in eukaryotic cells recovering from DNA damage. Here, we report a mechanism by which autophagy is suppressed in cells exposed to bacterial toxin-, chemical-, or radiation-mediated sources of genotoxicity. Autophagy suppression is directly linked to cellular responses to DNA damage, and specifically the stabilization of the tumor suppressor p53, which is both required and sufficient for regulating the ubiquitination and proteasome-dependent reduction in cellular pools of microtubule-associated protein 1 light chain 3 (LC3A/B), a key precursor of autophagosome biogenesis and maturation, in both epithelial cells and an *ex vivo* organoid model. Our data indicate that suppression of autophagy, through a newly identified p53-proteasome-LC3 axis, is a conserved cellular response to multiple sources of genotoxicity. Such a mechanism could potentially be important for realigning proteostasis in cells undergoing DNA damage repair.

## INTRODUCTION

Macroautophagy facilitates the degradation of normal cellular constituents under conditions of stress.^1^ During cellular recovery from DNA damage, the role of macroautophagy, which is critical for shaping and refining cytosolic proteostasis, is complex and poorly understood. Indeed, macroautophagy, which will hereafter be referred to interchangeably as “autophagy,” has been reported to be either suppressed^2–4^ or activated^5,6^ in a context-dependent manner during a cell’s response to DNA damage. Within DNA damaged cells, cytoprotective roles for autophagy have been proposed, including the degradation of pro-apoptotic proteins, removal of damaged organelles, and, the recycling of cellular goods to generate the dNTP building blocks and energy required for DNA repair.^7^ At the same time, substantial evidence exists that autophagy impairs cellular efforts to re-establish genome integrity and maintain cellular viability by degrading both anti-apoptotic and DNA damage response (DDR)-related proteins.^8,9^ Dissecting regulatory crosstalk between DDR and autophagy is often confounded by environmental conditions, some of which accompany select mechanisms of DNA damage, such as oxidative stress, that destabilize both genome integrity and proteostasis.^10–12^

To more directly evaluate the specific effects of DNA strand breaks on autophagy, we investigated cells that had been exposed to cytolethal distending toxins (CDTs), which are a family of conserved but broadly distributed protein genotoxins generated by a diverse group of bacteria that infect mucocutaneous barriers.^13^ CDTs generate DNA strand breaks within the nucleus of eukaryotic cells, but in the absence of additional off-target consequences associated with some DNA damaging agents that can activate cellular pathways not directly associated with DNA strand breaks.^14,15^ Here, our studies show that autophagy is suppressed in cells exposed to CDTs produced by several different pathogenic bacteria, as well as chemical- or radiation-induced sources of DNA damage. Mechanistically, autophagy was found to be negatively regulated by a mechanism involving the stabilization of the tumor suppressor protein p53, which promotes the ubiquitination and proteasome-mediated degradation of microtubule-associated protein 1 light chain 3B (LC3B), a critical component of autophagosome elongation and maturation.^16^ These studies reveal that suppression of macroautophagy, involving this newly identified p53-proteasome-LC3 axis, is a conserved cellular response to multiple types of DNA damage.

## RESULTS

### Autophagosome number is reduced in response to the DNA-damaging cytolethal distending toxin from *Campylobacter jejuni*

Macroautophagy is highly conserved among all eukaryotes as a cellular mechanism to degrade, recycle, and repurpose cytosolic contents under conditions of stress.^17^ For cells that have undergone the stress of DNA damage, the role of autophagy is not clear, as autophagy has been reported to be either suppressed^2^ or activativated^5,6^ in a context-dependent manner in cells exposed to DNA damaging agents. To readdress cellular regulation of autophagy in response to DNA strand breaks, human-derived HCT116 colonic epithelial cells, which are frequently employed as an *in vitro* model for studying DNA damage responses,^18,19^ were exposed to the *Campylobacter jejuni* cytolethal distending toxin (*Cj*-CDT). Toxin-dependent effects on autophagy were assessed by monitoring microtubule-associated protein 1 light chain 3B (LC3B), which is the most extensively studied of six human orthologs of the yeast autophagy protein Atg8.^20,21^ LC3B (which will hereafter be referred to as LC3) participates in several distinct stages within the multi-step process of autophagy, including autophagophore biogenesis and maturation, resulting ultimately in fully-formed autophagosomal vesicles that have engulfed cytosolic contents destined for recycling through lysosomal-mediated degradation.^16,20,21^ Immediately after expression, the carboxyl terminus of LC3 is processed, resulting in LC3-I,^22^ which is distributed diffusely within the cytosol. Under conditions of cellular stress such as nutrient starvation, LC3-I is conjugated with phosphatidylethanolamine, thereby generating LC3-II at nascent membranes of the developing phagophore.^20,23^

Immunofluorescence microscopy^24^ revealed that cells incubated with *Cj*-CDT possess visibly fewer and smaller LC3-enriched vesicles than untreated cells (Figure 1A), indicating that basal autophagy had been suppressed upon exposure to toxin. *Cj*-CDT-dependent activation of H2AX (γH2AX) was also observed, consistent with toxin-mediated DNA damage (Figure 1B).^25–27^ Immunoblot analysis revealed that cellular levels of LC3-II, which is widely used as a surrogate for autophagosome formation,^28^ were also significantly reduced in monolayers that had been incubated with *Cj-*CDT (Figure 1C). Taken together, these results are consistent with a conclusion that basal autophagy is suppressed in cells exposed to *Cj*-CDT.

**Figure 1.**
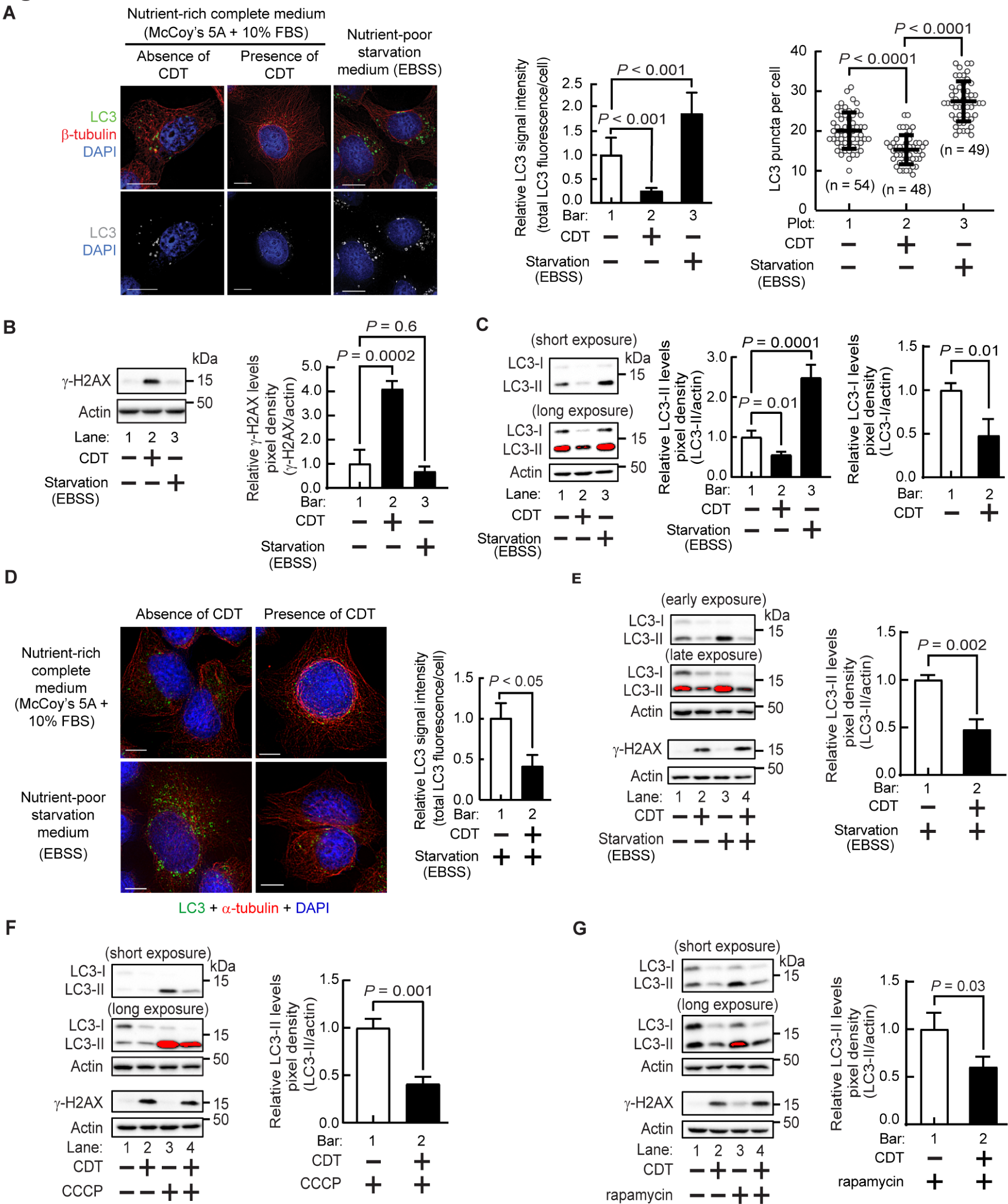
Effects of the DNA-damaging cytolethal distending toxin (*Cj*-CDT) on autophagy. HCT116 cells were incubated in McCoy’s 5A + 10% FBS at 37°C and under 5% CO_2_ in the absence or presence of *Cj*-CDT (10 nM). After 24 h, cell monolayers were further incubated with nutrient-rich McCoy’s 5A + 10% FBS (**A** to **G**), autophagy inducing nutrient-poor starvation medium (EBSS, Earl’s Balanced Salt Solution) (**A** to **E**), CCCP (25 μM) in McCoy’s 5A + 10% FBS (**F**), or rapamycin (20 μM) in McCoy’s 5A + 10% FBS (**G**). After 4 h, cells were imaged using fluorescence microscopy for total LC3-associated fluorescence signal intensity and total LC3 puncta (**A, D**), or, analyzed by immunoblot analysis for relative levels of LC3-II, γ-H2AX, and β-actin **(B** and **C, E** to **G)**. White scale bars from representative images indicate 10 μm (**A, D**). Images are representative of those collected from three biologically independent experiments (n=3) collected at 40X magnification. (**A, D**). Densitometric analyses of immunoblots (**B** and **C, E** to **G**) collected from 3 biologically independent experiments (n=3) were combined. Error bars represent standard deviations. Statistical analyses of the data were conducted using one-way Anova, followed by Dunnett’s post hoc test (**A** to **C**), or a two-tailed, unpaired Student’s *t*-test (**D** to **G**). Data are presented as mean ± SD. *P* < 0.05 indicating statistical significance (α = 0.05).

To assess whether *Cj*-CDT is sufficient to also suppress stress-mediated upregulation of autophagy, HCT116 cells that had been exposed to *Cj*-CDT were incubated for an additional 4 h under nutrient stress (EBSS medium). Both immunofluorescence imaging and immunoblot analysis revealed that cellular autophagy had been suppressed in a toxin-dependent manner (Figures 1D and 1E). Similar results were obtained in cells exposed to the widely-used alternative activators of autophagy, carbonyl cyanide m-chlorophenylhydrazone (CCCP) and rapamycin (Figures 1F and 1G), which activate macroautophagy by distinct mechanisms.^29,30^ Notably, we found no evidence that exposure to CCCP resulted in the activation of a specific form of autophagy called mitophagy, that selectively sequesters and degrades damaged mitochondria.^30,31^ Because both basal and activated autophagy were inhibited upon exposure to *Cj*-CDT (Figures 1A, 1C to 1G), our results indicate that DNA damage response-dependent LC3 degradation occurs either in the absence or presence of autophagy stimulation, but does not depend on the activation of autophagy.

### *Cj*-CDT-dependent reduction in cellular autophagosomes is not dependent upon enhanced lysosomal degradation

The results described above revealed that *Cj*-CDT-mediated DNA damage results in reduction of autophagosome number, but did not address the underlying mechanism. One possible explanation for the change in autophagosome number is an alteration in overall autophagic flux. Autophagic flux refers to the flow of cargo through the autophagic pathway, encompassing formation and expansion of phagophores into autophagosomes, which ultimately fuse with lysosomes to degrade sequestered contents.^29,32^ Steady-state levels of autophagosomes are shaped by processes that impact autophagosome biogenesis and turnover.^33^ A possible mechanism underlying *Cj*-CDT-dependent-reduction in cellular autophagosomes could involve an enhanced rate of autophagolysosomal degradation. To address this possibility, we examined the impact of *Cj*-CDT on autophagic flux in the presence or absence of inhibitors that block the degradative capacity of autophagolysosomes. These studies revealed that inhibition of lysosomal degradation did not prevent *Cj*-CDT-dependent reduction in the number of LC3-enriched autophagosomes (Figure 2A). In addition, the presence of lysosomal protease inhibitors did not prevent toxin-mediated reduction in cellular LC3-II levels (Figure 2B).

**Figure 2.**
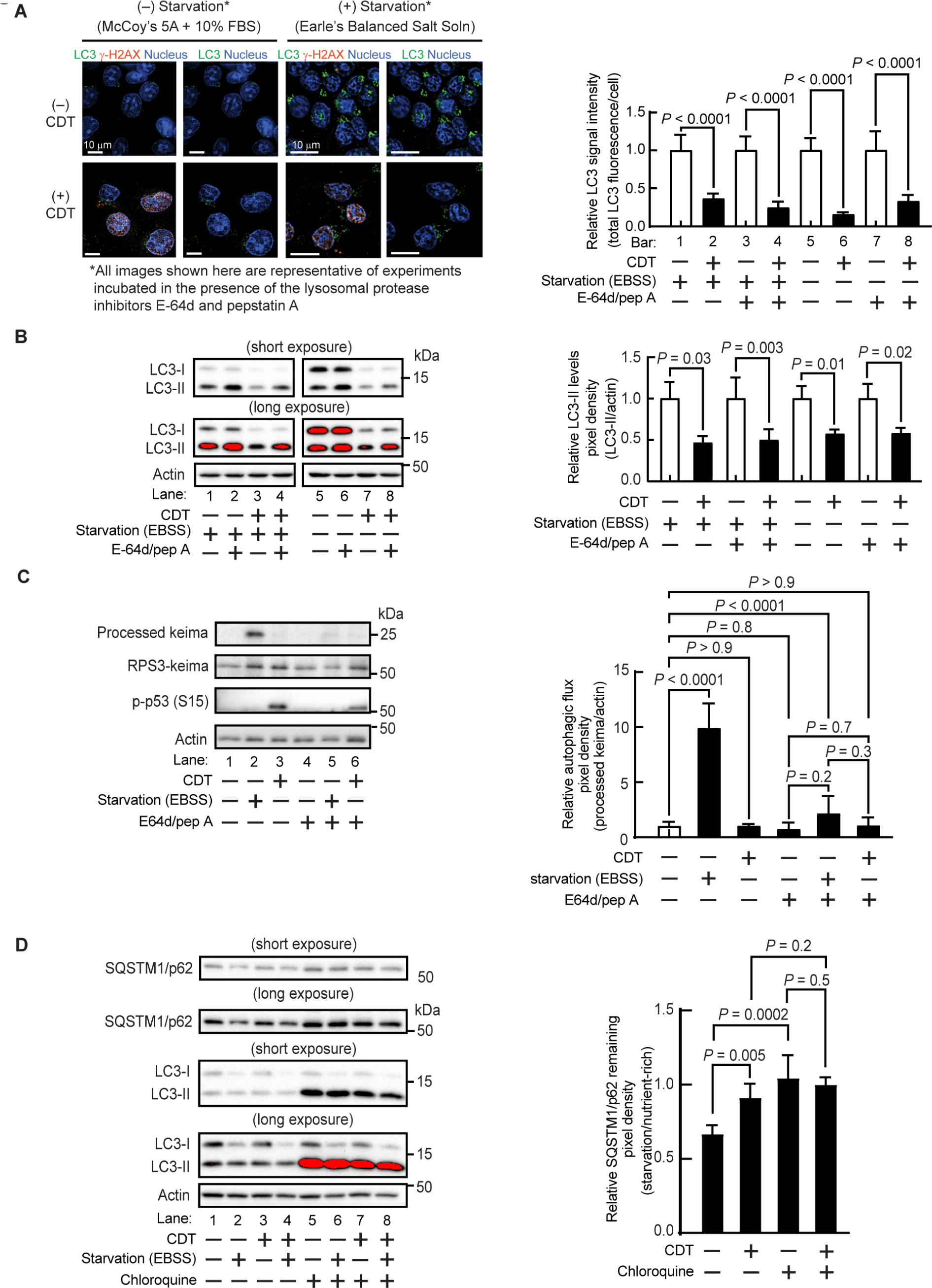
*Cj*-CDT-dependent effects on autophagolysosomal turnover. HCT116 cells within nutrient-rich McCoy’s 5A + 10% FBS medium, were incubated at 37°C in the absence (PBS pH 7.4) or presence of *Cj*-CDT (10 nM). After 24 h, the cell monolayers were washed 3 times with PBS. The monolayers were then further incubated at 37 °C and under 5% CO_2_, in the absence of toxin, with either EBSS or McCoy’s 5A + 10% FBS, as well as in the absence or presence of pepstatin A (10 μg/mL) and E-64d (10 μg/mL) (**A** to **C**), or, in the absence or presence of chloroquine (60 μM) and in the presence of actinomycin D (4 μM) (**D**). Relative levels of cellular LC3 were examined by fluorescence microscopy (**A**). White scale bars in representative images indicate 10 μm (**A**). Relative levels of LC3-II (**B**), mKeima (**C**), and SQSTM/p62 (**D**) were examined by immunoblot analysis. Images collected at 40X magnification (**A**) and immunoblots (**B** to **D**) are representative of those collected from three biologically independent experiments (n=3). Data from 3 biologically independent experiments (n=3) were combined and are shown (**A** to **D**). Error bars represent standard deviations of data combined from 3 biologically independent experiments (n=3). Statistical analyses of the data were conducted using unpaired, two-tailed, Student’s t-test (**A** and **B**), or, two-way Anova, followed by Fisher’s LSD post hoc test (**C** and **D**). Data are presented as mean ± SD. *P* < 0.05 indicates statistical significance (α = 0.05).

The results above are consistent with a conclusion that CDT does not upregulate autophagic flux. To more directly validate this conclusion, we employed a recently-developed approach for monitoring degradation of bulk autophagic cargo, which involves stable expression of the protease-sensitive ribosomal protein S3 (RPS3) fused to the amino-terminus of the hyperstable protein mKeima.^34^ *Cj*-CDT-dependent alterations in autophagic flux were evaluated by assessing changes in the electrophoretic mobility of mKeima, resulting from autophagolysosomal degradation of RPS3-mKeima (55 kDa) into the lower molecular weight species comprising mKeima only (referred to as “processed mKeima”, 25 kDa). As expected, cellular levels of processed mKeima were significantly greater in cells where autophagy had been activated by nutrient starvation (Figure 2C), consistent with a profound upregulation of autophagic flux. In contrast, under conditions where *Cj*-CDT-dependent stabilization of p53 (phospho-p53 (Ser15)) is observed, the cellular levels of processed mKeima were nearly undetectable, and visually indistinguishable, in the lysates of cells incubated in either the absence or presence of toxin (Figure 2C).

We extended the studies described immediately above by evaluating the association between increased lysosomal degradative activity and enhanced autophagic flux in cells that had been exposed to *Cj*-CDT. Specifically, we evaluated the degradation of SQSTM1/p62, an adaptor protein that targets cargo to the autophagosome,^32^ in cells that had been incubated in the absence or presence of toxin. As expected, monolayers of cells incubated in the absence of *Cj*-CDT exhibited a reduction in cellular SQSTM1/p62 levels under starvation conditions relative to nutrient-rich conditions, consistent with autophagy-mediated degradation of SQSTM1/p62. In contrast, there was less starvation-induced reduction of SQSTM1/p62 in monolayers incubated in the presence of *Cj*-CDT, similar to cells incubated in the presence of the autophagic flux inhibitor chloroquine (Figure 2D), which impairs lysosome fusion with autophagosomes.^35^ Taken together, these experiments support a model that *Cj*-CDT-dependent reduction in autophagosome number is not likely dependent upon upregulation of lysosomal degradative activity within the autophagic pathway.

### *Cj*-CDT-mediated inhibition of autophagy occurs downstream of autophagic activation

We next evaluated the possibility that *Cj-*CDT-mediated suppression of autophagy involves modulation of cellular signaling that regulates autophagy. An important regulator of autophagy activation under conditions of stress is mTORC1 (the mechanistic target of rapamycin complex 1). Activation of mTORC1 is characterized by increased anabolism and protein synthesis, while inhibition of mTORC1 is characterized by upregulation catabolism and activation of autophagy.^29^ A commonly used marker of mTORC1 activity, p70 S6 kinase (S6K), is directly phosphorylated by mTORC1 at threonine 389 (p-S6K(Thr389)), leading to an increase in protein synthesis.^29,36^ Under conditions of cellular stress, mTORC1 is inhibited, leading to a loss of p-S6K(Thr389).

Studies to determine if *Cj*-CDT-dependent inhibition of autophagy results from inhibition of mTORC1 activity revealed that p-S6K(Thr389) levels were approximately the same in cells that had been incubated in the absence or presence of *Cj*-CDT (Figure S1A), indicating that toxin-dependent suppression of autophagy is not associated with inhibition of mTORC1. To further validate these results, we next assessed a potential role for an essential upstream mediator of autophagosome biogenesis, which is the Unc-51-like autophagy activating kinase 1 (ULK1) complex.^37^ Normally, the ULK1 complex is negatively regulated by mTORC1 through phosphorylation at serine 757 (p-ULK1 (Ser757)).^38^ Consistent with our finding that *Cj*-CDT-dependent suppression of autophagy does not involve toxin modulation of mTORC1 signaling, immunoblot analyses also revealed that activation of the ULK1 protein kinase complex was approximately the same in cells that had been incubated in either the presence or absence of *Cj*-CDT (Figure S1B). Collectively, our results thus far are consistent with a model of *Cj*-CDT-dependent suppression of autophagy that involves alterations in a critical step that occurs downstream of the signaling required for autophagy activation, but, prior to lysosomal-mediated turnover of autophagosome contents.

### *Cj*-CDT-dependent suppression of autophagy is linked to depletion of available pools of cellular LC3

Because autophagosome biogenesis requires the conversion of available LC3-I into the lipid conjugated LC3-II,^28^ we next evaluated the possibility that *Cj*-CDT-dependent DNA damage may be linked to altered cellular availability of LC3. Immunoblot analyses of lysates from HCT116 cells that had been exposed to *Cj*-CDT revealed that, under conditions resulting in H2AX activation (γ-H2AX), cellular levels of LC3-I were visibly and quantitatively reduced (Figure 1C), supporting the idea that *Cj*-CDT-dependent suppression of autophagy may be associated with available cellular pools of LC3.

Experiments to identify specific Atg8 family members^39^ involved in *Cj*-CDT-dependent suppression of autophagy revealed that only cellular levels of LC3A and LC3B (Figures S1C and S1D), but not LC3C, GABARAP, GABARAPL1, or GABARAPL2 (Figures S1E to S1H), were reduced in HCT116 cells that had been exposed to *Cj*-CDT. These results suggest that *Cj*-CDT-dependent suppression of autophagy may entail the selective involvement of LC3B/A.

### *Cj*-CDT-dependent reduction of cellular LC3 was not idiosyncratic to HCT116 cells, as reduction in cellular LC3 was recapitulated across several unrelated cell lines

Cellular LC3-I levels were significantly reduced by approximately 55% in human embryonic kidney HEK293-T cells exposed to *Cj*-CDT (10 nM), and by approximately 50% in murine fibroblast NIH/3T3 cells (Figures S2A and S2B). In addition, *ex vivo* experiments demonstrated that LC3-I was significantly reduced in mouse intestinal organoids (approximately 40%) that had been exposed to *Cj*-CDT (Figure S2C).

Significant reductions in cellular LC3 were also observed when HCT116 cells were incubated with CDTs from either *Escherichia coli* (*Ec*-CDT), or *Haemophilus ducreyi* (*Hd*-CDT) (Figures S2D and S2E). Although our sampling was not exhaustive, these results suggest that the capacity to induce reductions in cellular LC3 may be shared among individual members of the CDT genotoxin family.

*Cj*-CDT-dependent depletion of both LC3-I and LC3-II was detected after 24 h at a toxin concentration of 10 nM, but not at 1 nM (Figures S3A and S3B), similar to the concentrations of *Cj*-CDT required for DNA damage in HCT116 cells (Figure 1B), which is in accordance with the premise that toxin activity was responsible for the cellular reduction of LC3. Additional experiments revealed that LC3-I reduction was detected after 18 h, but not 12 h of exposure to *Cj*-CDT (10 nM). In comparison, reduced LC3-II cellular levels were detected after 24 h but not after 18 h (Figure S3C). These results are consistent with a model for autophagy suppression involving toxin-mediated reduction in available LC3, which by hindering autophagosome biogenesis, impairs the ability of cells to fully replace the number of mature LC3-enriched autophagosomes that are degraded during normal autophagic flux.

### *Cj*-CDT-dependent DNA damage precedes and is required for the depletion of cellular LC3

We next more closely assessed the relationship between *Cj*-CDT-mediated DNA damage and the reduction in cellular LC3 levels within monolayers exposed to the toxin. The active domain of *Cj*-CDT, *Cj*-CdtB, possesses DNase I-like catalytic activity that results in single- and double-strand DNA breaks in the nucleus of host cells, and the subsequent activation of the DDR.^27,40,41^ To evaluate a possible causal linkage between the DNA strand-breaking activity of *Cj*-CDT, and, toxin-dependent LC3 depletion, HCT116 cells were exposed to purified wild-type *Cj*-CDT or purified mutant *Cj*-CDT (CdtB (H157G)) (Figure S4A), whose catalytic domain is deficient in DNase I activity^42^ but is still capable of localization to the nucleus within intoxicated cells (Figure S4B). These experiments validated the requirement for the DNA strand-breaking activity of *Cj*-CdtB for both the reduction in cellular LC3 levels, and, the suppression of autophagy (Figures S4C and S4D).

Additional studies revealed that *Cj*-CDT-dependent DNA damage, as indicated by elevated levels of cellular γ-H2AX,^25,26^ was detected after 12 h of toxin exposure but not after 1 h (Figure S4E). Because depletion of LC3-I levels was detected at 18 h but not at 12 h (Figure S3C), our results indicate that *Cj*-CDT-dependent DNA damage precedes the depletion of cellular LC3, which is consistent with the idea that the reduction in LC3-enriched vesicles may occur as a consequence of toxin-mediated genotoxicity.

### Suppression of autophagy is a conserved response to multiple types of DNA damage

To evaluate the possibility that suppression of autophagy is a conserved cellular response to different mechanisms of DNA damage, and not limited to single- and double-stranded DNA breaks in cells exposed to the CDTs, cells were incubated with the well-characterized DNA damaging agents Bleocin, which causes the generation of reactive free radicals,^43^ etoposide, which is a topoisomerase-II inhibitor,^44^ 5-fluorouracil (5-FU), which causes deoxynucleotide pool imbalances,^14^ and UV irradiation, which causes formation of thymine dimers.^14,45,46^ These experiments revealed that, similar to studies described above for cells incubated with *Cj*-CDT (Figure 1C), reduced levels of LC3 coincided with increased levels of γ-H2AX in cells that had been exposed to Bleocin, etoposide, 5-FU, or UV radiation (Figure S5). In the course of these studies, we did not observe measurable cell death at concentrations of each DNA damaging agent associated with activation of DDR and reduction in cellular LC3 levels (Figure S6), ruling out a scenario that a reduction in viable cells was responsible for the suppression of the cellular autophagic response observed throughout our studies. These results suggest that DDR-dependent suppression of autophagy is a conserved and widely-distributed response to multiple, alternative mechanisms of DNA damage.

### Reduction in LC3-I or LC3-II was not observed within DNA-damaged cells overexpressing LC3B

Because our data suggested that *Cj*-CDT-dependent suppression of autophagy is a consequence of depleted cellular LC3 pools, we predicted that ectopic overexpression of LC3B, which is rapidly converted to LC3-I, should obviate DNA damage-dependent suppression of autophagosome biogenesis. Experiments to test this prediction revealed that *Cj*-CDT- or Bleocin-dependent decreases in LC3-I or LC3-II were not observed within HCT116 cells overexpressing LC3B (Figures 3A and 3B). Control experiments indicated that ectopic overexpression of LC3B did not interfere with *Cj*-CDT- or Bleocin-mediated activation of H2AX (Figure 3C). These data suggest that, within DNA damaged cells, impairment in autophagy upregulation is due directly to insufficient cellular pools of LC3-I available for conversion into LC3-II as an obligate prerequisite for autophagosome biogenesis.

**Figure 3.**
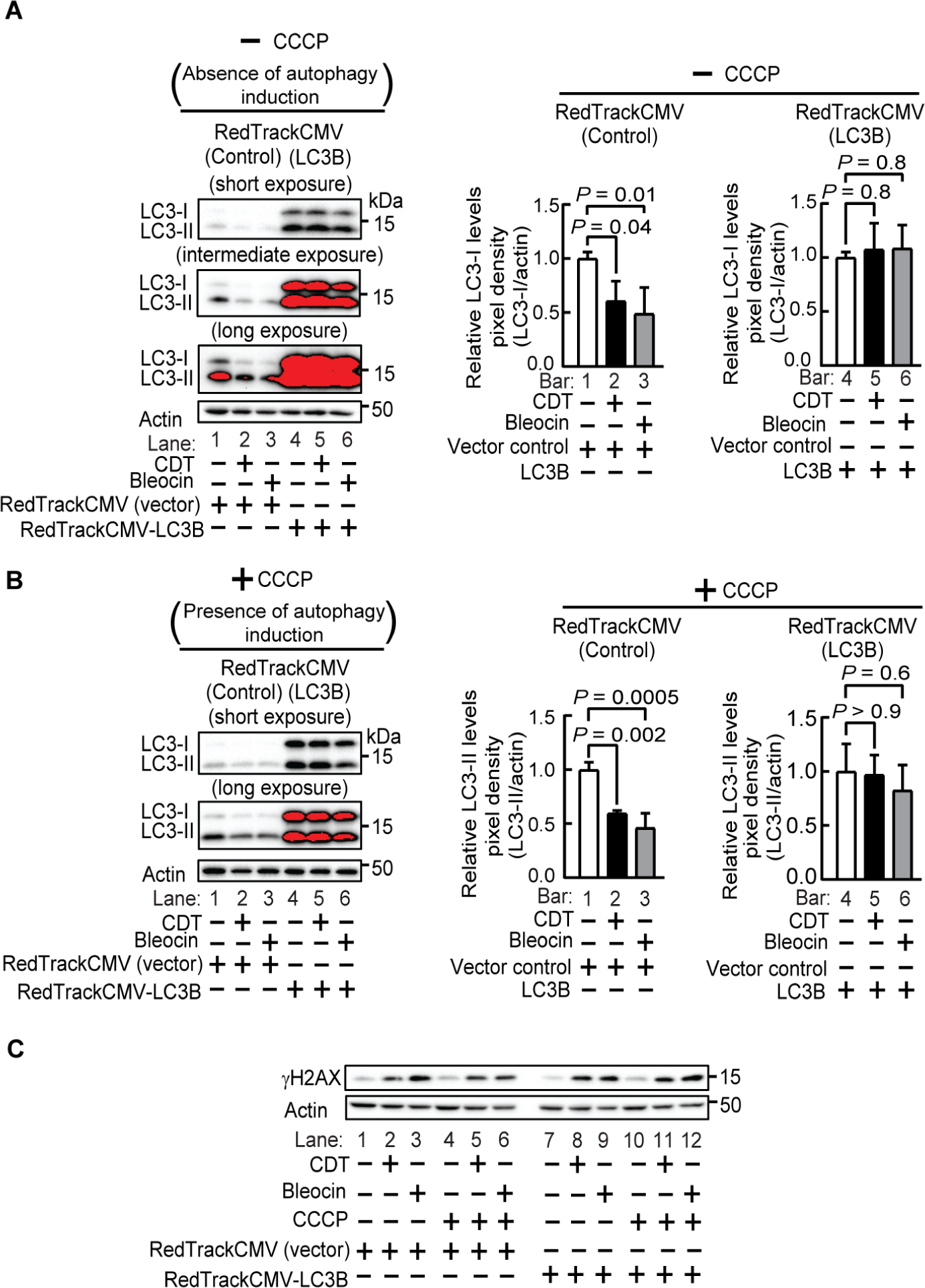
Depletion of LC3-I is causal for the reduction in cellular LC3-II in cells exposed to *Cj*-CDT. HCT116 cells transfected with RedTrackCMV or RedTrackCMV-LC3B were incubated with *Cj*-CDT (10 nM) or Bleocin (10 μg/mL) (**A** to **C**). After 24 h, cell lysates were further incubated for 3 h in the absence (**A** and **C**) or presence (**B** and **C**) of the autophagy inducer, CCCP (25 μM), and then analyzed by immunoblot analysis for LC3-I and LC3-II (**A** and **B**), or, γ-H2AX (**C**). Densitometric analyses of immunoblots collected from 3 biologically independent experiments (n=3) were combined. Error bars represent standard deviations. Statistical analyses of the data were conducted using one-way Anova, followed by Dunnett’s post hoc test (**A** and **B**). Data are presented as mean ± SD. *P* < 0.05 indicates statistical significance (α = 0.05).

### ATM-dependent DNA damage response (DDR) is causally associated with reduction in basal LC3 cellular levels

Cells exposed to *Cj*-CDT exhibit the capacity to mobilize a robust DNA damage response (DDR)^27,47–49^ which is coordinated, in part, by the DNA damage signaling kinase, ataxia telangiectasia mutated (ATM).^50^ To evaluate a potential association between the responsiveness of ATM to DNA damage, and, *Cj-*CDT-dependent depletion of LC3 levels, we examined HCT116 cells exposed to *Cj-*CDT (10 nM) for 24 h in the presence of KU-55933, a potent inhibitor of ATM kinase activity.^50^ These studies demonstrated that, in HCT116 monolayers exposed to *Cj*-CDT, and under conditions of both basal and up-regulated autophagy, inhibition of ATM kinase activity with KU-55933 was sufficient to rescue levels of cellular LC3 (Figure 4A), thereby supporting a causative link between ATM-dependent DDR and depletion of LC3 in toxin-treated cells.

**Figure 4.**
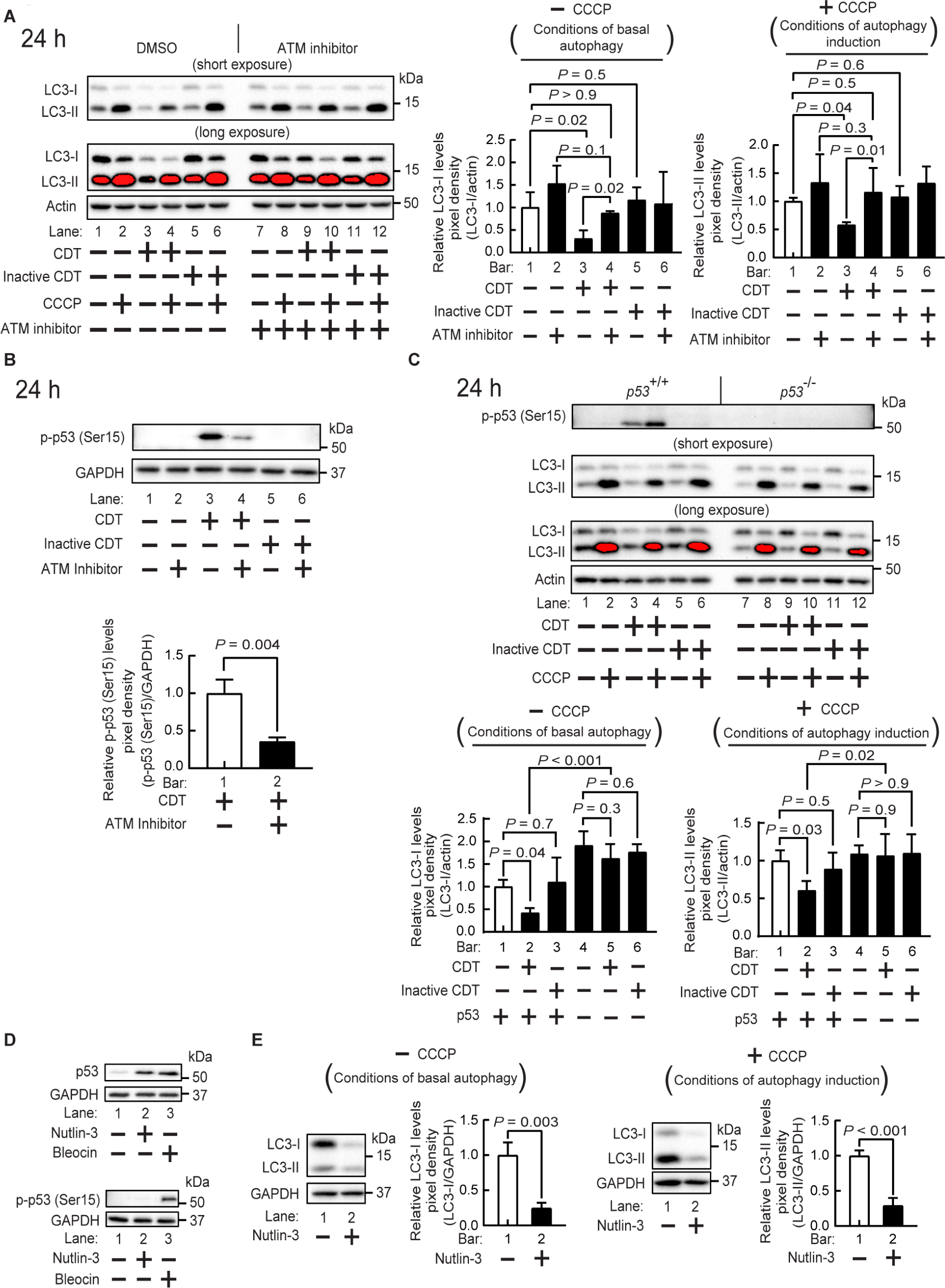
The DNA damage response is important for CDT-dependent reduction in both cellular LC3-I and LC3-II levels. HCT116 (*p53*^+/+^) cells (**A** to **E**) or HCT116 (*p53*^-/-^) cells (**C**) were preincubated with KU-55933 (10 μM) (**A** and **B**), and then in the absence or presence of *Cj*-CDT (10 nM) (**A** to **C**), inactive CDT (containing *Cj*-CdtB H157G, 10 nM) (**A** to **C**), or Bleocin (10 μg/mL) (**D**). Alternatively, cell monolayers were incubated in the absence or presence of Nutlin-3 (20 μM) (**D** and **E**). After 24 h, monolayers were further incubated with CCCP (25 μM) (**A, C,** and **E**). Lysates were analyzed by immunoblot analysis for relative levels of LC3-I (**A, C,** and **E**), LC3-II (**A, C,** and **E**), β-actin (**A** and **C**), total p53 (**D**) p-p53 (S15) (**B** to **D**), and GAPDH (**B, D,** and **E**). The immunoblots are representative of those from 3 independent experiments (n=3). Densitometric analyses were combined from 3 independent experiments (n=3), and the relative levels of LC3-I, LC3-II, and p-p53 were calculated by dividing the intensity of immuno-specific bands corresponding to cellular LC3-I (**A, C,** and **E**), LC3-II (**A, C,** and **E**), and p-p53 (**B**) by the intensity of immuno-specific bands corresponding to controls β-actin (**A** and **C**) or GAPDH (**B** and **E**). The data represented by the black bars were each separately rendered relative to the data represented by the white bars, which were assigned an arbitrary value of 1.0. Error bars represent standard deviations. Two-way Anova followed by Fisher’s LSD post hoc test (**A** and **C**), or two-tailed, unpaired Student’s *t*-test (**B** and **E**) were used. Data are presented as mean ± SD. *P* < 0.05 indicates statistical significance (α = 0.05).

### Within DNA damaged cells, stabilization of p53 is essential and sufficient for suppression of autophagy due to depletion of cytosolic LC3

Among potential ATM-regulated effector proteins associated with DDR, the tumor suppressor protein p53 was earlier reported to be activated by ATM within CDT intoxicated cells.^48,51^ HCT116 cells possess genetically wild-type p53 at both alleles, which is stabilized upon irradiation.^52^ We exposed HCT116 cells to *Cj*-CDT, and confirmed that p53^48,51^ is stabilized by phosphorylation at serine-15 (p-p53 (Ser15)) (Figure 4B), and that p-p53 (Ser15) generation was inhibited in the presence of the ATM inhibitor, KU-55933 (10 μM) (Figure 4B). In addition, LC3-I levels were no longer reduced, and, autophagy was no longer suppressed in response to *Cj*-CDT in cells lacking p53 (HCT116 *p53* ^-/-^) (Figure 4C). These results support a model that stabilization of p53 is required for suppression of autophagy within DNA damaged cells.

Because p53 stabilization can occur by mechanisms extending beyond DDR,^52–55^ we next evaluated the hypothesis that p53 stabilization alone is sufficient to suppress autophagy. For these studies, degradation of cellular p53, in the absence of DNA damaging agents, was prevented by using Nutlin-3, an inhibitor of MDM2, the E3-ubiquitin ligase responsible for targeting p53 for degradation.^56^ These studies revealed that, as expected, cellular p53 was readily detected within HCT116 cells that had been exposed to Nutlin-3 (Figure 4D). In addition, p-p53 (Ser15) was not detected in Nutlin-3 treated cells (Figure 4D), indicating that stabilization had occurred in the absence of ATM-mediated phosphorylation. Importantly, under these conditions, both depletion of cellular LC3 and suppression of autophagy were clearly evident (Figure 4E), similar to that observed in DNA damaged cells. These results indicate that stabilization of cellular p53 is a key step in the mechanism by which LC3 levels are reduced, ultimately resulting in the suppression of autophagy.

### *Cj*-CDT-dependent reduction in cellular LC3 levels occurs by an autophagy-independent mechanism

Having established a clear link between DNA damage and LC3 reduction, we next turned our attention to address the mechanism by which LC3 is degraded in DNA damaged cells. It is known that LC3-II, which is incorporated into autophagosome membranes by a process requiring the Atg5-Atg12 conjugation complex^57^, is normally degraded through autophagolysosomal turnover.^32^ Our data thus far (Figure 2) suggested that increased autophagosome turnover is not likely the predominant mechanism by which overall cellular LC3 is reduced in CDT-treated cells. However, these same data do not rule out the possibility that autophagic degradation of LC3 may nonetheless contribute to the overall decrease in cellular LC3 following exposure to *Cj*-CDT. Studies designed to directly address this possibility revealed *Cj*-CDT-dependent reduction of cellular LC3-I in both Atg5^+/+^ and Atg5^-/-^ cells which, respectively, are able or unable to facilitate conjugation of LC3-I to PE as a required step for autophagosome biogenesis (Figure 5A).^32,58^ These results indicate that autophagy is not essential for *Cj*-CDT-dependent reduction in cellular LC3, thereby supporting a model of DNA damage-dependent LC3 reduction that involves a non-autophagic degradative pathway.

**Figure 5.**
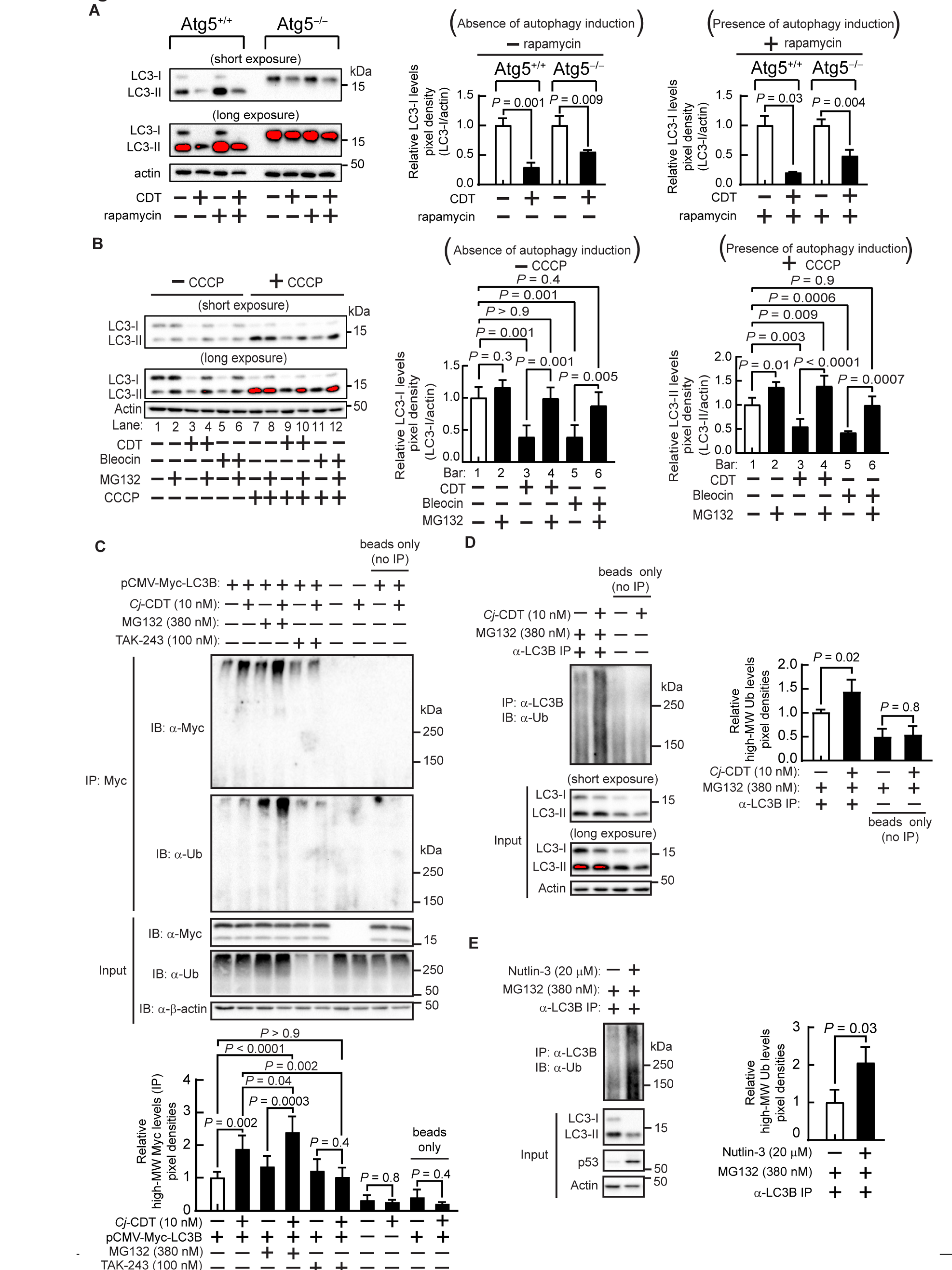
DNA damage-dependent suppression of autophagy is linked to the ubiquitination and depletion of LC3 by a proteasome-dependent mechanism. MEF Atg5^+/+^ and MEF Atg5^-/-^ cells were incubated in the absence or presence of *Cj*-CDT (10 nM) (**A**). After 48 h, MEF cells were further incubated with rapamycin (20 μM) (**A**). After 3 h in the presence of rapamycin, cell lysates were examined for LC3-I, LC3-II, and β-actin (**A**). HCT116 cells (**B** to **E**), either transfected (**C**), or, not transfected (**B** to **E**) with the pCMV-Myc-LC3 overexpression plasmid, were incubated with *Cj*-CDT (10 nM) (**B** to **D**), Bleocin (10 μg/mL) (**B**), or Nutlin 3 (20 μM) (**E**), and, in the presence or absence of MG132 (380 nM) (**B** to **D**) or TAK-243 (100 nM) (**C**). After 24 h, HCT116 monolayers were further incubated with CCCP (25 μM) (**B**), while Nutlin-treated monolayers were further incubated with MG132 (380 nM) (**E**). After 3 h in the presence (**B**) or absence (**B** to **D**) of CCCP, or 4 h in the presence of MG132 (**E**), cell lysates were examined for LC3-I (**A** to **E**), LC3-II (**A** to **E**), Myc-LC3B (**C**), ubiquitin (**C**), total p53 (**E**), and β-actin (**B** to **E**). Alternatively, Myc-LC3B was immunoprecipitated with Myc-specific antibodies, and ubiquitinated Myc-LC3B was detected using Myc-specific and ubiquitin-specific antibodies (**C**). Intrinsic cellular LC3B was immunoprecipitated using LC3-specific antibodies,and ubiquitinated LC3B was detected using ubiquitin-specific antibodies (**D** and **E**). Immunoblots are representative from data collected from at least 3 biologically independent experiments (n=3). Densitometric analyses were combined from three independent experiments. The data shown by the black (filled) bars were each separately rendered relative to the data represented by the white (empty) bar, which was assigned an arbitrary value of 1.0. Error bars represent standard deviations. Statistical analyses of the data were conducted using two-tailed, unpaired Student’s *t*-test (**A** and **E**) or two-way Anova, followed by Fisher’s LSD post hoc test (**B** to **D**). Data are presented as mean ± SD. *P* < 0.05 indicates statistical significance (α = 0.05).

### Within DNA damaged cells, LC3 is depleted by a proteasome-dependent mechanism

Based on results described above indicating that DNA-damage dependent LC3 reduction does not require the involvement of an autophagic degradative pathway, we next addressed potential alternative mechanisms by which cellular pools of LC3 are reduced in response to DNA damage. Studies to examine the mechanism by which LC3 pools in DNA damaged cells are depleted failed to provide evidence that either cellular levels of *MAP1LC3B* transcripts (Figure S7A), or, overall protein synthesis (Figure S7B), were altered in cells exposed to *Cj*-CDT. However, *Cj-*CDT-dependent reduction in basal steady state levels of cellular LC3 was inhibited in cells exposed to the proteasome inhibitor, MG132 (Figure 5B), which suggested that within DNA damaged cells, LC3 is depleted by a proteasome-dependent mechanism. Similar results were observed in HCT116 cells incubated with the DNA damaging agent Bleocin (Figure 5B), indicating that proteasome-dependent depletion of cellular LC3 is recapitulated in response to distinct sources of DNA damage.

### Within DNA damaged cells, cellular LC3 is ubiquitinated

The results described above indicate that cellular proteasome activity is important for *Cj*-CDT-dependent depletion of LC3. Since protein degradation via the proteasome system can occur via either ubiquitination-dependent or -independent mechanisms of proteasome substrates,^59,60^ we next evaluated whether LC3 is ubiquitinated within DNA damaged cells. Within monolayers exposed to *Cj*-CDT, immunoblot analyses revealed a visible and quantitative increase in high molecular weight ubiquitinated forms of ectopically overexpressed Myc-LC3 (Figure 5C). Moreover, an increase in high molecular weight, ubiquitinated LC3 species was observed in lysates of cells in which proteasome degradative activity was inhibited with MG132 (Figure 5C), further supporting the importance of the proteasome for depletion of available LC3 in cells exposed to *Cj*-CDT. At the same time, *Cj-*CDT-dependent production of high molecular weight LC3 species was reduced in the presence of the ubiquitin-activating enzyme inhibitor TAK-243 (Figure 5C), congruent with a model that post-translational modification with ubiquitin is important for targeting LC3 for proteasome-mediated degradation. Notably, a ladder of bands, corresponding to the addition of individual, 8 kDa ubiquitin monomers to a growing polyubiquitin chain, was not detected in these studies, consistent with previous observations associated with complex ubiquitination patterns comprising multiple linkage types and branching.^61^ In addition, we also detected visible and quantitative increases in high molecular weight ubiquitinated species in *Cj*-CDT-treated cells corresponding to endogenous LC3 (i.e. in the absence of ectopic expression of LC3) (Figure 5D), consistent with an interpretation that LC3 ubiquitination was not an artifact of ectopic Myc-LC3 overexpression. Immunoblot analyses of the LC3-specific immunoprecipitates did not reveal the presence of the abundant autophagophore / autophagosome-associated protein p62, suggesting that ubquitinated forms of LC3-phagophore associated proteins are not responsible for the signal in our IP experiments (Figure S8). Taken together, these results support a model of DNA-damage-dependent suppression of autophagy that occurs by a mechanism involving proteosome degradation of ubiquitinated LC3. Notably, unrelated studies previously identified both monoubiquitinated and polyubiquitinated forms of LC3B that are degraded at the proteasome.^62–66^

### Stabilization of p53 is sufficient for ubiquitination and proteasome degradation of LC3

The experiments described thus far indicate that Cj-CDT-dependent suppression of autophagy requires the ubiquitination and proteosome-mediated degradation of LC3.

In addition, our data indicate that stabilization of p53 is required for reduction in cellular LC3 pools. In further support of a causative link between stabilization of p53 and proteosome-mediated degradation of ubiquitinated-LC3, immunoblot analysis revealed significantly higher levels of ubiquitinated-LC3 from lysates of cells that had been exposed to Nutlin-3 (Figure 5E), which, as described above, effectively stabilizes p53 in the absence of DNA damage. These results indicate that stabilization of cellular p53 is a key step for the suppression of autophagy by a mechanism requiring the ubiquitination and proteasome degradation of LC3.

### DDR-dependent suppression of autophagy is reversible

The suppression of autophagy within DNA damaged cells is potentially an important component of a cell’s “emergency response” to maintain viability while reestablishing genome integrity. However, because long-term impairment of autophagy would likely be detrimental to cellular health, restoration of functional autophagy is likely to be a critical component for reconstituting overall cellular homeostasis. To evaluate the plasticity by which autophagy is regulated within DNA damaged cells, HCT116 monolayers were exposed to either *Cj-*CDT or Bleocin. After 24 h, at conditions under which LC3 is depleted and autophagy is suppressed, inhibition of ATM kinase activity resulted in a significant reduction in cellular p53 stabilization (Figure 6A) and, notably, restoration of overall cellular LC3 levels (Figure 6B), consistent with the conclusion that DDR-dependent suppression of autophagy is reversible. These studies support a model that p53 functions as a molecular rheostat for regulating autophagy under conditions of DNA damage, thereby allowing cellular repair responses to be relatively nimble as conditions change.

**Figure 6.**
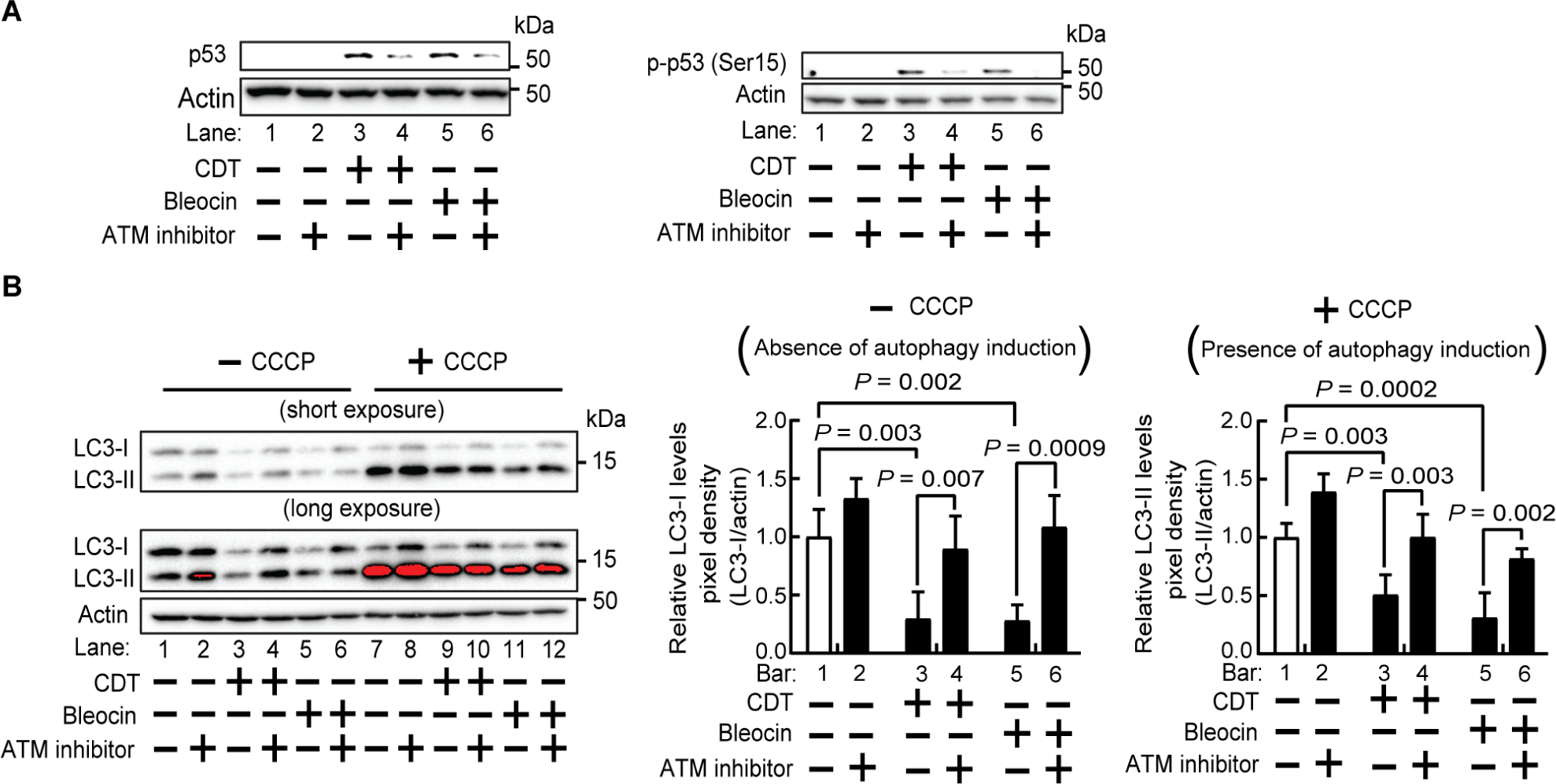
DNA damage-dependent reduction in both cellular LC3-I and LC3-II levels is reversed by impairing ATM-mediated stabilization of p53. HCT116 cells were incubated for 24 h with *Cj*-CDT (10 nM), or Bleocin (10 μg/mL). After 24 h, monolayers were further incubated in the absence or presence of the autophagy inducing compound CCCP (25 μM) (**B**), as well as the absence or presence of the ATM inhibitor, KU-55933 (10 μM). After an additional 3 h, cell lysates were examined, using immunoblot analyses, for total p53 (**A**), p-p53 (Ser15) (**A**), LC3-I (**B**), LC3-II (**B**), and β-actin (**A** and **B**). The immunoblots shown are representative from 3 biologically independent experiments (n=3). Densitometric analyses were combined from 3 immunoblots (**B**), and the relative levels of cellular LC3-I or LC3-II levels were calculated. The data shown by the black (filled) bars were each separately rendered relative to the data represented by the white (empty) bar, which was assigned an arbitrary value of 1.0. Error bars represent standard deviations of data combined from 3 biologically independent experiments (n = 3). Statistical analyses of the data were conducted using two-way Anova, followed by Fisher’s LSD post hoc test. Data are presented as mean ± SD. *P* < 0.05 indicates statistical significance (α = 0.05).

## DISCUSSION

Here we report that autophagy is suppressed within DNA damaged cells through a newly identified p53-proteasome-LC3 axis (Figure 7). This model is supported by several principal findings. First, autophagy is negatively regulated in cells exposed to a DNA damaging bacterial genotoxin (Figure 1 and Figure 2), as well as multiple chemical or radiation-mediated sources of DNA damage (Figure S5). Second, in cells experiencing DNA damage, autophagy is regulated through DDR (Figure 4), and in particular, the reversible modulation of p53 stability (Figure 6). Third, modulation of p53 stability regulates the ubiquitination and proteasome-mediated degradation of LC3, thereby impacting cellular pools of this autophagy effector (Figure 5). These results contribute to and are consistent with mounting evidence for interconnectivity between cellular DNA damage, DDR, and autophagy.^2,5,6,67,68^

**Figure 7.**
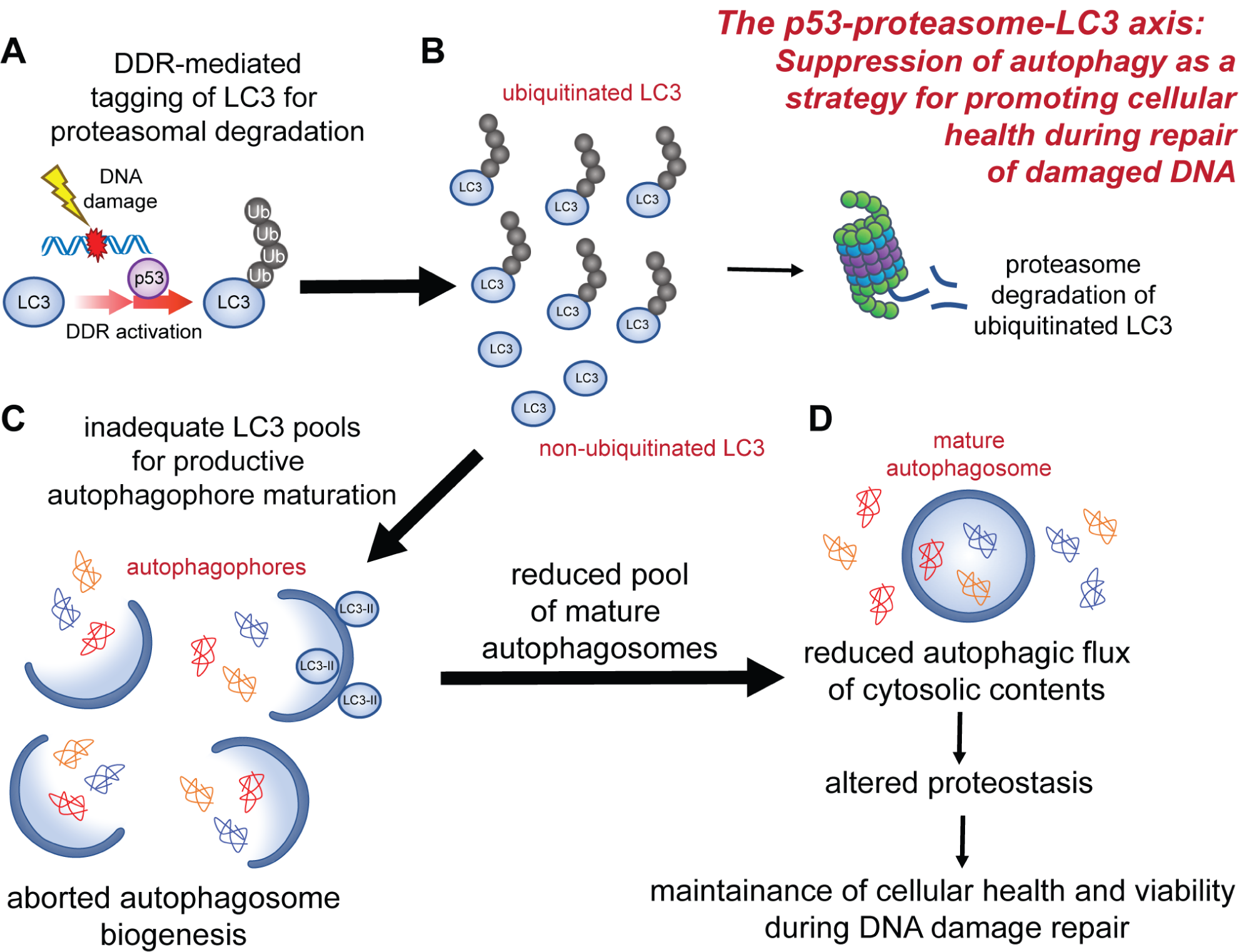
Model of the proposed p53-proteasome-LC3 axis for regulating the autophagic response in DNA damaged cells. Autophagy suppression results from proteasomal degradation of LC3 within DNA damaged cells as a mechanism for maintaining viability during restoration of genome integrity (**A** to **D**). Upon DNA damage, DDR activation leads to stabilization of p53, which is required for ubiquitination of LC3 (**A**), thereby targeting LC3 for proteasome degradation (**B**). Available LC3 pools for autophagosome maturation are diminished leading to the production of fewer mature autophagosomes (**C**). Fewer mature autophagosomes results in reduced autophagic flux and preservation of pro-survival factors (**D**).

Autophagy has been reported to be either suppressed^2–4,69,70^ or activated^5,6,67,71,72^ during a cell’s response to DNA damage, suggesting that regulation of autophagy through DDR is complex and occurs in a context dependent manner. For example, within DNA damaged cells, cytoprotective roles for autophagy have been proposed, including the degradation of pro-apoptotic proteins, removal of damaged mitochondria, and, the recycling of cellular goods to generate the dNTP building blocks and energy required for DNA repair. Several lines of evidence support an important role for the degradation of cellular factors within DNA damaged cells through autophagic flux. For example, autophagy-deficient (beclin1^-/-^) tumor cells were earlier found to accumulate DNA strand breaks,^71^ suggesting a functional role for increased autophagic flux in promoting cellular repair of DNA damage. Another example is autophagy-dependent reduction in cellular levels of HP1α, a chromatin-associated factor, which suggests a direct role in facilitating access of DNA repair proteins to sites of double stranded breaks^67,72^. At the same time, substantial evidence exists that autophagy impairs cellular efforts to re-establish genome integrity and maintain cellular viability by degrading both anti-apoptotic and DDR-related proteins. For example, autophagy has been reported to play a role in the degradation of the DNA damage repair protein, Sae2.^9^ Another example is that p53 was found to inhibit autophagy in irradiated lung cancer cells.^3^ Additionally, GADD45A, a p53-regulated DNA damage response protein, has been shown to inhibit autophagy.^4^ Because autophagy is one of several primary mechanisms (including transcription, translation, proteosome mediated degradation, etc.) responsible for managing cellular proteostasis, we speculate that context-dependent up- or down-regulation of autophagy in DNA damaged cells may provide the nimbleness needed for cells to pivot in response to shifting cellular priorities. Additional studies will be required to resolve the mechanisms underlying differential autophagy-regulation in response to DNA damage.

Our studies identified p53 stability as a critical determinant in regulating autophagy through proteasome-mediated degradation of ubiquitinated LC3. To our knowledge, this is the first demonstration that links cellular responses to DNA damage involving p53 to proteasomal-mediated regulation of LC3 as a mechanism for regulating autophagy. As noted above, our findings are supported by unrelated studies that recently identified both monoubiquitinated and polyubiquitinated forms of LC3B that are degraded at the proteasome.^62–66^ The exact role of p53 in proteasomal regulation of autophagy remains an important and intriguing question. Preliminary studies in our laboratory, using a non-exhaustive siRNA screen of several p53-regulated E3 ligases, failed to identify an E3 ligase responsible for LC3 ubiquitination in CDT-treated cells (not shown). p53-dependent down-regulation of LC3 mRNA was previously reported in chronically starved cells,^73^ suggesting a potential role for p53 as a transcriptional factor. However, neither a change in LC3 transcripts nor a reduction in overall protein synthesis in DNA-damaged cells was observed in our studies (Figure S7). Nonetheless, given the large transcriptome regulated by p53 as a transcription factor, we cannot rule out the possibility that p53 regulates the expression of intermediary factors that contribute to LC3 ubiquitination. In addition, non-transcriptional-related activities of p53 have also been described.^53,74^ For example, p53 directly interacts with and enhances ubiquitination of the multi-functional regulator, 14-3-3 gamma.^75^ Thus, it remains possible that, in response to DNA damage, p53 mediates LC3 ubiquitination by a mechanism independent of transcriptional regulation.

Although our results indicate that DNA damage-dependent suppression of autophagy is associated with a reduction in overall cellular LC3, depletion of this critical component could not be attributed solely to heightened degradative capacity of autophagolysosomes (Figure 2). Rather, our studies revealed DNA damage dependent ubiquitination and degradation of LC3. In addition to LC3A/B, all Atg8 homologs share a ubiquitin-like core structural motif consisting of a four-stranded β-sheet flanked by two α-helices. Although the reasons underlying the differential modulatory effects of DNA damage on cellular levels of Atg8 sub-family members are not presently understood, it is noteworthy that several recent, unrelated publications reported that LC3 subfamily members (LC3A, LC3B, and LC3C), but not members of the GABARAP subfamily, are subject to ubiquitination, and subsequent proteosome-mediated degradation.^63,65,66^ Moreover, ubiquitination of LC3A and LC3B was recently demonstrated to be reversible through the action of the deubiquitinase, USP10.^66^ The authors of these studies suggested that cycles of LC3B ubiquitination and deubiquitination impact autophagic activity by controlling cellular LC3B, although these authors did not address the mechanism by which these ubiquitination cycles might be regulated. In the case of cells that have experienced DNA damage, our data suggest a key role for p53 in regulating autophagy through modulation of proteasome-mediated turnover of LC3A/B. Currently, it is not clear to what extent p53-dependent modulation of autophagy in cells might extend beyond DNA damage to other cellular stresses.

The demonstration that autophagy is reversibly suppressed in DNA damaged cells, by a mechanism involving the DDR-dependent inhibition of autophagosome formation, is consistent with the idea that cellular proteostasis is altered in response to DNA damage. ^76,77^ By degrading specific cellular proteins in response to changing environmental demands, autophagy likely plays a critical role in both the maintenance and remodeling of cellular proteostasis. Although the full spectrum of changes in the cellular proteome that occurs in response to DNA damage is incompletely understood, preliminary studies in our laboratory found that cellular levels of CtIP,^78^ the mammalian ortholog of yeast Sae2, a DNA damage repair protein previously shown to be targeted by autophagy, were elevated in HCT116 cells under conditions in which we detect *Cj*-CDT-dependent autophagy suppression (Figures S9A and S9B). In contrast, cellular levels of CtIP were reduced in DNA-damaged cells ectopically overexpressing LC3B (Figures S9A and S9B). Congruent with the idea that, in the face of DNA damage, suppression of autophagy might promote cellular survival, we also found that apoptosis was elevated when LC3B was overexpressed in DNA damaged cells (Figures S9C and S9D). Although not exhaustive, results from these preliminary studies are congruent with the hypothesis that *Cj*-CDT-mediated autophagy suppression might confer a protective advantage to cells that have experienced DNA damage through the stabilization of cellular factors that promote the re-establishment of genome integrity.

In summary, this study illustrates a new mechanism, involving a newly identified p53-proteasome-LC3 axis, by which the suppression of autophagy in response to DNA damage may function as a major driver in realigning cellular proteostasis in cells undergoing DDR. Suppression of autophagy appears to be a conserved response to multiple sources of DNA damage. Delineating how independent but concurrent sources of cellular stress (e.g., DNA damaging agents and oxidative stress) impinge upon cellular decision making will be important for achieving a holistic view of how cells respond to the challenges of genotoxicity. Finally, from a clinical perspective, these studies suggest that augmentation of DNA-damaging cancer treatments with pro-autophagic drugs could potentially impair physiological DNA repair responses, and thus in some cases, increase the efficacy of existing treatment regimens.^79,80^

## Author Contributions

DJL, MKC, and SRB designed the majority of the experiments and wrote the manuscript. KAB and HP provided invaluable consultation and guidance regarding experimental design and made substantial contributions to the writing of the manuscript. MKC optimized and conducted experiments for Figure 1, Figure 2, Figure 3, Figure 4, Figure 5, Figure 7, Figure S1, Figure S2, Figure S5, Figure S6, Figure S7, Figure S8, Figure S9, and provided statistical consultation for all experiments. JRK optimized and conducted experiments for Figure 1, Figure 2, Figure 3, Figure 4, Figure S3, and Figure S5. BT optimized and conducted experiments for Figure 1, Figure 2, and Figure S4. PK optimized and conducted experiments for Figure S3 and Figure S5. IJK optimized and conducted experiments for Figure S9. YL contributed to experiments for Figure 3, Figure 5, Figure 6, and Figure S4. EJ optimized and conducted experiments for Figure S1 and Figure S2. ME contributed to experiments for Figure 1, Figure 4, and Figure S4. BT collected and analyzed immunofluorescent microscopy images for Figure 1 and Figure 2. HC contributed to development and implementation of quality control measures for CDT production. SBS and HM contributed to the development and optimization of recombinant CDT production. All authors reviewed the manuscript.

## Acknowledgements

We acknowledge the Flow Cytometry Facility (CMtO) at the Roy J. Carver Biotechnology Center at the University of Illinois at Urbana-Champaign (specifically Barbara Balhan and Dr. Barbara Pilas for determining cell cycle progression), Dr. Bert Vogelstein (Johns Hopkins Medicine) for the HCT116 p53 WT/KO cell lines, Elizabeth Good (University of Illinois) for the NIH/3T3 cell line, Dr. Noboru Mizushima (Tokyo Medical and Dental University) for the Mouse embryonic fibroblast (MEF) Atg5^-/-^ and MEF Atg5^+/+^ cell lines, Dr. Heeseon An (Sloan Kettering Institute at Memorial Sloan Kettering Cancer Center) for the HCT116 RPS3-Keima cell line, and the Kuzminov Lab (University of Illinois) for the assistance and use of their Hoefer UV-C 500 Crosslinker. SYBR Green PCR master mix was a gift from Dr. Jie Chen (University of Illinois) and prepared by Nilmani Singh (Jie Chen Lab, University of Illinois).

## MATERIALS AND METHODS

### RESOURCE AVAILABILITY

#### Lead Contact

Further information and requests for resources and reagents should be directed to and will be fulfilled by the lead contact, Dr. Steven Blanke

#### Materials Availability

Plasmids generated in this study are available from the lead contact.

#### Data and Code Availability

All data reported in this paper (microscopy images, original western blot images, original protein gel images, raw flow cytometry data, raw microplate reader data, and raw RT-qPCR data) will be shared by the lead contact upon request. This paper does not report original code. Any additional information required to reanalyze the data reported in this paper is available from the lead contact upon request.

### EXPERIMENTAL MODEL AND SUBJECT DETAILS

#### Mice

All animal work was reviewed and approved by the Illinois Institutional Animal Care and Use Committee (IACUC protocol #15238). Murine intestinal organoids were harvested from female C57BL/6 mice purchased from Jackson Laboratories. 20 – 25 week old mice were used for generation of organoid culture.

#### Cell Culture

All mammalian cells were maintained at 37°C in a humidified atmosphere under CO_2_ (5%). HCT116 (human colon cancer epithelial cell line, male; CCL-247, ATCC, USA) cells, including HCT116 (p53^+/+^, a gift from Dr. Bert Vogelstein, Johns Hopkins Medicine), HCT116 (p53^-/-^, a gift from Dr. Bert Vogelstein), and HCT116 RPS3-Keima (stably expressing mKeima fused to the C-terminus of RPS3, a gift from Dr. Heeseon An) were maintained in McCoy’s 5A (modified) medium with L-Glutamine (#SH30200.01, GE Healthcare, USA) medium supplemented with a 10% final concentration of fetal bovine serum (FBS; #12306C, Lot. 16C315, Sigma-Aldrich, USA). HEK293T (human embryonic kidney epithelial cell line, female; CRL-3216, ATCC, USA) cells were maintained in Dulbecco’s Modified Eagle’s Medium (DMEM; #10-013-CV, Corning, USA) supplemented with a 10% final concentration of FBS. NIH/3T3 (Mouse embryonic fibroblast cell line, male, a gift from Elizabeth Good, University of Illinois Urbana-Champaign) cells were maintained in DMEM (#10-013-CV, Corning, USA) supplemented with a 10% final concentration of FBS. MEF (mouse embryonic fibroblast) (Atg5^-/-^, a gift from Dr. Noboru Mizushima) and MEF (Atg5^+/+^, a gift from Dr. Noboru Mizushima) were maintained in DMEM (#10-013-CV, Corning, USA) supplemented with a 10% final concentration of FBS. Cell lines were tested for mycoplasma contamination using MycoAlert Mycoplasma Detection Kit (LT07-118, Lonza Biosciences, Switzerland).

#### Murine Intestinal Organoids

Murine intestinal organoids were harvested from the small intestine of 20 – 25 week old female C57BL/6 mice. Cultured 3D organoids were embedded in Matrigel (#354234, Corning, USA) and maintained in IntestiCult Mouse Organoid Growth Medium (#06005, Stem Cell Technologies, Canada). Organoids were incubated at 37°C in a humidified atmosphere under CO_2_ (5%).

#### Bacterial Strains

*Escherichia coli* BL21 used for protein expression and purification were grown at 37 °C in Luria broth (LB) supplemented with ampicillin (100 μg/mL final concentration) to select for *E. coli* BL21 containing the desired pET-15b expression plasmid. *E. coli* DH5α used for plasmid purification were grown in LB supplemented with ampicillin (100 μg/mL final concentration, for pCMV-myc-LC3) or kanamycin (50 μg/mL final concentration, for RedTrackCMV and RedTrackCMV-LC3B).

### METHODS DETAILS

#### Expression and purification of recombinant CDTs

Recombinant CDT subunits (CdtA, CdtB, and CdtC) derived from *Campylobacter jejuni* (*Cj*-CDT), *Haemophilus ducreyi* (*Hd*-CDT), and *Escherichia coli* (*Ec*-CDT) were expressed and purified separately, as described previously.^81^ Briefly, the genes encoding the individual CdtA, CdtB, and CdtC subunits (within pET-15b protein expression plasmids; #69661, Sigma-Aldrich, USA), each expressed with an amino-terminal polyhistidine fusion peptide, were expressed separately in *E*. *coli* BL21 (DE3) protein expression strains. 1 L of Luria broth (LB) in 6 L flasks were inoculated with 1-3 mL starter cultures of *E*. *coli* BL21 (DE3) and were cultivated at 37 °C with rotary agitation. Once cultures reached an optical density at 600 nm (OD_600_) of 0.4-0.6, cultures were further cultivated with rotary agitation at 37 °C in the presence of 0.2 mM isopropyl1-thio-β-D-galactopyranoside (IPTG) to induce CDT subunit expression. After 3 h, cultures were harvested by centrifugation at 8,000 *× g* at 4 °C. After 8 min, pellets were resuspended in 10 mL of ice-cold PBS pH 7.4 and disrupted via sonication (Ampl. 25%, 2 sec ON / 5 sec OFF, Total ON = 3 min; Model FB505 (500 Watt), Fisher Scientific, USA) on ice and in the presence of 1 mM phenylmethylsulfonyl fluoride (PMSF). After sonication, contents were centrifuged at 8,000 *× g* at 4 °C. After 30 min, the supernatant was discarded and pellets resuspended in 15 mL of 8 M urea solution (8 M urea, 20 mM HEPES, 200 mM NaCl in water, pH 7.5 at 4°C) and agitated on a rotary shaker on ice. After 30 min, contents were disrupted on ice via sonication (Ampl. 25%, 2 sec ON / 5 sec OFF, Total ON = 3 min; Model FB505 (500 Watt), Fisher Scientific, USA). The sonicated cell lysates were centrifuged at 8,000 *× g* at 4 °C. After 30 min, the supernatant was transferred into clean conical tubes and purified by cobalt metal affinity chromatography using approximately 4-6 mL of cobalt resin slurry (Talon Metal Affinity Resin, #635503, Clontech Laboratories, USA) within a gravity-flow column. Columns were washed with 1 mM imidazole (1 mM imidazole in 8 M urea solution at pH 7.5) and eluted with 100 mM imidazole (100 mM imidazole in 8 M urea solution at pH 7.5) for the CdtA subunit and approximately 25 mM imidazole (25 mM imidazole in 8 M urea solution at pH 7.5) for the CdtB and CdtC subunits. The concentrations of each of the purified subunits were then determined using a BCA Protein Assay Kit (#23225, Thermo Fisher Scientific, USA). Eluted subunits were then diluted to 100 μg/mL with 8 M urea solution, combined in equimolar amounts of CdtA, CdtB, and CdtC subunits in a final volume of around 200 mL, and then transferred into regenerated cellulose dialysis tubing (6000 – 8000 MWCO; #21-152-4, Fisher Scientific, USA). Equimolar amounts of CdtA, CdtB, and CdtC within 8 M urea solution were dialyzed, on a stir plate at 4 °C, against 4 L of HEPES buffer (20 mM HEPES, 200 mM NaCl, and 5% glycerol in water, pH 7.5). After every 6 h, for the next 12 h, dialysis tubing was transferred into fresh 4 L of HEPES buffer and dialyzed on a stir plate at 4 °C. After 18 h of dialysis in HEPES buffer, dialysis tubing containing CDT was transferred into 4 L of phosphate buffer (PB buffer, 0.456g NaH_2_PO_4_, 4.343g Na_2_HPO_4_·7H_2_O, 5% glycerol in 1 L H_2_O, pH 7.5) and dialyzed on a stir plate at 4 °C. After 6 h, dialysis tubing was transferred into fresh 4 L of PB buffer and incubated on a stir plate at 4 °C. After 12 h of dialysis in PB buffer, for *Cj*-CDT holotoxin, contents of the dialysis tubing, around 200 mL, were concentrated by binding the diluted CDT holotoxin from the dialysis tubing onto a gravity flow anion exchange column (DEAE SephaceI, #17050001, GE Healthcare, USA) containing 3 mL of slurry, and then eluted with 10 mL of PBS pH 7.4. Approximately, 500 μL fractions were collected and analyzed for the presence of all three subunits as well as increased protein band intensity using SDS-PAGE and Coomassie Brilliant Blue staining. For *Ec*- and *Hd*-CDT holotoxins, contents of the dialysis tubing were transferred into centrifugal filters (Amicon Ultra, 10,000 MWCO; #UFC901024, Sigma-Aldrich, USA) in order to concentrate the diluted holotoxin post-dialysis. To isolate holotoxin containing equimolar concentrations of CdtA, CdtB, and CdtC subunits, concentrated fractions containing CDT holotoxin were loaded onto a gravity flow size exclusion column (1 g, P-60 resin; #1504164, Bio-Rad, USA) pre-equilibrated with PBS pH 7.4 as the mobile phase and 500 μL elution fractions taken.

Fractions were then analyzed for stoichiometric equivalency of CdtA, CdtB, and CdtC subunits by assessing for approximately equal band intensities of the CdtA, CdtB, and CdtC subunits using SDS-PAGE and Coomassie Brilliant Blue staining. Fractions found to contain approximately stoichiometrically equivalent amounts of CdtA, CdtB, and CdtC subunits were aliquoted into 10 μL aliquots, with each aliquot containing toxin in PBS pH 7.4, and stored at −80 °C. The concentration of the purified holotoxin was determined using a BCA Protein Assay Kit prior to storage. Activity of purified CDTs was assessed through cell cycle phase assays, as described below.

#### Cell cycle phase determination

For each preparation of CDT used in these studies, we calculated a cell cycle arrest dose_50_ (CCAD_50_ value), which defines the concentration of holotoxin required to induce arrest of 50% of the specific cell type within the monolayer. The indicated cell lines were seeded (approximately 0.1 × 10^6^ cells per well) on 12-well culture plates and incubated at 37 °C and under 5% CO_2_. After 24-36 h, the cells were incubated at 37 °C and under 5% CO_2_ in complete medium containing CDT at the indicated concentrations, or, were incubated with PBS pH 7.4 in place of CDT. After 24 h, cell culture medium containing CDT or PBS pH 7.4, as the carrier control, was removed and cells were washed twice with PBS pH 7.4. Cells were then detached from the cell culture plate by trypsinization at 37 °C. After detachment of cells was observed, 10% final concentration of FBS in PBS pH 7.4 was added to inhibit trypsin activity. Cells were then pelleted (500 *× g*, 5 min) and resuspended by adding ice cold 70% ethanol dropwise with constant vortexing. Cells were incubated at −20 °C in the presence of 70% ethanol. After a minimum of 1 h, cells were pelleted (800 *× g*, 5 min) and resuspended in PBS pH 7.4.

After 30 min at room temperature, cells were pelleted and resuspended in staining solution (300 μL per sample; containing 0.1% Triton X-100, 1 mg/mL RNase A, and 10 μg/mL propidium iodide in PBS pH 7.4). After 1 h at room temperature, cells were analyzed using a BD FACS Canto II (BD Biosciences, USA) flow cytometry analyzer. Flow cytometry data were processed using FCS Express software (De Novo Software, USA) and statistics performed using GraphPad Prism 7 (GraphPad Software, USA).

#### Induction of autophagy through nutrient starvation

Cellular monolayers that were incubated at 37 °C and under 5% CO2 in nutrient-rich complete medium were washed three times with PBS pH 7.4 in order to remove residual nutrient-rich medium. Cellular monolayers were then incubated at 37 °C and under 5% CO_2_ in Earl’s Balanced Salt Solution (EBSS; #24010043, Thermo Fisher Scientific, USA) for up to 8 h. For non-starved controls, cellular monolayers were instead incubated at 37 °C and under 5% CO2 for up to 8 h in nutrient-rich complete medium.

#### DIC/Fluorescence Microscopy

8-well cell culture chamber slides (Lab-Tek II; #154534, Thermo Fisher Scientific, USA) were analyzed using a DeltaVision RT microscope (GE Life Sciences, USA). Olympus Plan Apo x60 and x100 oil objectives were used. Images were acquired using a Photometrics CoolSnap HQ camera (Photometrics, USA). Images were processed using SoftworX Explorer Suite (Version 3.5.1, GE Life Sciences, USA). For each cell, images were collected from an average of approximately 30 Z-planes using 0.2 μm intervals. Deconvolution was performed using SoftworX constrained iterative deconvolution tool (ratio mode) and analyzed using Imaris (Version 7.4.2, Bitplane, Switzerland).

#### Assessing autophagosome levels

Cells were seeded (approximately 20,000 cells per well) on 8-well cell culture microscope slides and incubated at 37 °C and under 5% CO_2_. After 24 – 36 h, the cells were incubated at 37 °C and under 5% CO_2_ according to treatment described in each individual experiment (see figure legends). At the completion of the experiment, cell monolayers were washed three times with PBS pH 7.4. Cell monolayers were then fixed with 100% ice cold methanol. After 15 min at −20 °C, fixative was removed, and cell monolayers washed three times with PBS pH 7.4. Then, cells were permeabilized with 0.1% Triton X-100 in PBS pH 7.4 at room temperature. After 15 min, cell monolayers were then blocked with blocking buffer (5% final concentration of bovine serum albumin in PBS, pH 7.4). After 1 h, cell monolayers were washed three times with PBS-T (PBS, pH 7.4, containing 0.1% final concentration of Tween-20), 5 min per wash. Cell monolayers were incubated at 4 °C with rabbit monoclonal anti-LC3A/B antibodies (#12741, Cell Signaling Technology, USA) at a 1:100 dilution in PBS-T. Additional primary antibodies against β-tubulin (1:50 dilution in PBS-T; #86298, Cell Signaling Technology, USA) and/or γ-H2AX (1:200 dilution in PBS-T; #9718, Cell Signaling Technology, USA) were added as indicated (see figure legends). After overnight primary antibody incubation, cell monolayers were washed three times with PBS-T, 5 min per wash. Cell monolayers were then incubated in the presence of secondary antibody (goat anti-rabbit IgG (H+L) Alexa Fluor 488 conjugate (#A11008, Thermo Fisher Scientific, USA), or, donkey anti-mouse IgG (H+L) Alexa Fluor conjugate 568 (#A10037, Thermo Fisher Scientific, USA), 1:1000 dilution in PBS-T). After 1 h at room temperature, cell monolayers were washed three times with PBS-T, 5 min per wash.

DAPI (500 ng/mL) staining solution was then added to the cellular monolayers. After 30 min at room temperature, cell monolayers were washed three times with PBS-T, 5 min per wash. Prolong Gold antifade reagent (#P36930, Thermo Fisher Scientific, USA) was then added to stained monolayers and a coverslip was applied and sealed with clear nail polish. Images were acquired as described above. LC3 fluorescence signal intensity per cell was quantified by dividing the LC3 fluorescence signal intensity within the entire field by the cell number within the field. This value was then corrected for the increase in the average size of cells undergoing *Cj*-CDT-mediated cell cycle arrest (which is approximately 1.7 ± 0.2 times larger in two-dimensional area than cells incubated with PBS, pH 7.4, as the carrier control). LC3 puncta per cell were quantified by manually counting visible puncta of LC3 fluorescence signal for each whole cell within the field.

For each biological replicate, 6 - 10 microscopic fields at 40X magnification, each containing approximately 20-30 cells, were analyzed.

#### Lysate preparation and Immunoblotting

The indicated cell lines were seeded (approximately 0.04-0.05 × 10^6^ cells per well) on 24-well culture plates and incubated at 37 °C and under 5% CO_2_. After 24 – 36 h, the cells were incubated according to the treatment described in each individual experiment (see figure legends). At the indicated time of harvest, 24-well plates were placed on ice and cell monolayers were washed twice with cold PBS pH 7.4. After washing the monolayers, 75-150 μL of RIPA buffer (#89901, Thermo Fisher Scientific, USA) supplemented with HALT protease and phosphatase inhibitor cocktail (#78443, Thermo

Fisher Scientific, USA) were added to each well and the plate incubated at 4 °C on a platform rocker with occasional agitation. After 20 min, lysates were transferred to chilled microcentrifuge tubes and Laemmli sample buffer (6X; #LB0100, Morganville Scientific, USA), supplemented with β-mercaptoethanol, was added to generate a 1X concentration of Laemmli sample buffer in cell lysate. Lysates, on ice, were then passed through a 25-gauge needle three times to fully homogenize samples and then boiled at 100°C for 10 min. Samples were fractionated by sodium dodecyl sulfate-polyacrylamide vertical gel electrophoresis using a range of 4-20% polyacrylamide gels and transferred to PVDF membranes using a wet/tank blotting system. Membranes were blocked at room temperature with 5% non-fat milk in TBS-T (0.1% Tween-20 in TBS pH 8.0) or 5% bovine serum albumin in TBS-T, as indicated in the relevant methods section. After 1 h, membranes were washed with TBS-T to remove residual blocking buffer and then incubated at 4 °C with the indicated primary antibodies diluted in TBS-T. After an overnight incubation with primary antibodies, membranes were washed with TBS-T, three times, 10 min per wash, and membranes were subsequently incubated at room temperature with horseradish peroxidase (HRP)-conjugated secondary antibodies diluted in TBS-T. After 1 h, membranes were washed with TBS-T, three times, 10 min per wash, and then HRP-conjugated secondary antibody binding visualized using enhanced chemiluminescent substrate for HRP (1:5 mixture of SuperSignal West Femto Maximum Sensitivity Substrate (#24095) to SuperSignal West Pico Plus Chemiluminescent Substrate (#34580); Thermo Fisher Scientific, USA). Immunoblots were imaged using the ChemiDoc system (XRS+, Bio-Rad, USA). Immunoblot densitometry analyses were performed using Image Lab software (Version 4.1, Bio-Rad, USA).

#### Immunoblot densitometric analyses

Image Lab image documents (.scn) were opened in Image Lab software (Version 4.1, Bio-Rad, USA). Lanes of the immunoblot images were denoted, and bands detected using the Image Lab software. Band detection area was adjusted to ensure that any background signal was not included in the quantification. Relative quantities of target proteins were determined by dividing the protein band volumes (as calculated from pixel densities) of the target proteins by the protein band volumes of the reference proteins [loading controls used: β-actin, GAPDH, or α-tubulin]. Statistics were performed using GraphPad Prism 7 (GraphPad Software, USA). The specific statistical test used is noted within each figure legend. For each biological replicate, statistical analyses and comparisons of data that had undergone densitometric quantification were conducted using data derived from a single immunoblot.

#### Assessing DNA damage response initiation

Cellular lysates were fractionated by SDS-PAGE using 12% polyacrylamide gels, transferred to PVDF membranes, and membranes were blocked with 5% non-fat milk in TBS-T. Membranes were then incubated with rabbit monoclonal anti-phospho-histone H2AX (Ser139) (γ-H2AX) (#9718, Cell Signaling Technology, USA) antibodies at a 1:3000 dilution in TBS-T, followed by anti-rabbit IgG, HRP-linked secondary antibodies (#7074, Cell Signaling Technology, USA) at a 1:10,000 dilution in TBS-T. HRP-conjugated secondary antibody binding, immunoblot imaging, and immunoblot densitometry analysis were performed as described above.

#### Assessment of cellular levels of LC3-II and LC3-I

Cellular lysates were fractionated by SDS-PAGE using 12% polyacrylamide gels, transferred to PVDF membranes, and membranes were blocked with 5% non-fat milk in TBS-T. Membranes were then incubated with rabbit monoclonal anti-LC3A/B antibodies (#12741, Cell Signaling Technology, USA) at a 1:3000 dilution in TBS-T, followed by anti-rabbit IgG, HRP-linked secondary antibodies (#7074, Cell Signaling Technology, USA) at a 1:10,000 dilution in TBS-T. HRP-conjugated secondary antibody binding, immunoblot imaging, and immunoblot densitometry analysis were performed as described above.

#### Autophagy induction using CCCP

Cell culture medium was removed from cellular monolayers that were incubated at 37 °C and under 5% CO_2_. Cellular monolayers were then incubated at 37 °C and under 5% CO_2_ in cell culture medium containing carbonyl cyanide m-chlorophenyl hydrazine (CCCP, 25 μM final concentration; #C2759, Sigma-Aldrich, USA) for up to 3 h. For the non-autophagy inducing control treatment, cellular monolayers were instead incubated at 37 °C and under 5% CO_2_ in cell culture medium containing the carrier control dimethyl sulfoxide (DMSO, 0.1% final concentration; #D2650, Sigma-Aldrich, USA), for up to 3 h. After 3 h, the cell monolayers were harvested, and lysates were prepared for immunoblot analyses as described above in “Lysate preparation and Immunoblotting”.

#### Autophagy induction using Rapamycin

Cell culture medium was removed from cellular monolayers that were incubated at 37°C and under 5% CO_2_. Cellular monolayers were then incubated at 37 °C and under 5% CO_2_ in cell culture medium containing rapamycin (20 μM final concentration; #S1039, SelleckChem, Germany) for up to 4 h. For the non-autophagy inducing control treatment, cellular monolayers were instead incubated at 37 °C and under 5% CO_2_ in cell culture medium containing the carrier control DMSO (0.1% final concentration) for up to 4 h. After 4 h, the cell monolayers were harvested, and lysates prepared as described previously.

#### Inhibition of lysosomal turnover

Cell culture medium was removed from cellular monolayers that were incubated at 37 °C and under 5% CO_2_. Cellular monolayers were then incubated at 37 °C and under 5% CO_2_ in cell culture medium containing the lysosomal protease inhibitors E-64d (10 μg/mL final concentration; #E8640, Sigma-Aldrich, USA) and pepstatin A (10 μg/mL final concentration; #P5318, Sigma-Aldrich, USA) for up to 4 h. Cells treated with the carrier control were instead incubated at 37 °C and under 5% CO_2_ in cell culture medium containing DMSO (0.2% final concentration) for up to 4 h.

#### Assessment of RPS3-Keima processing

HCT116 cells stably expressing RPS3-Keima were seeded (approximately 0.04-0.05 × 10^6^ cells per well) on 24-well culture plates and incubated at 37 °C and under 5% CO_2_. After 24 – 36 h, cell culture medium was removed and cells were further incubated in the absence or presence of *Cj*-CDT (10 nM) under nutrient-rich conditions (McCoy’s 5A + 10% FBS), or in nutrient deplete medium (EBSS), in either the absence or presence of the lysosomal protease inhibitors E64d (10 μg/mL final concentration; #E8640, Sigma-Aldrich, USA) and pepstatin A (10 μg/mL final concentration; #P5318, Sigma-Aldrich, USA). After 24 h, monolayers were washed and approximately 100 μL of RIPA buffer (#89901, Thermo Fisher Scientific, USA) supplemented with HALT protease and phosphatase inhibitor cocktail (#78443, Thermo Fisher Scientific, USA) and Pierce Universal Nuclease (#88702, Thermo Fisher Scientific, USA) were added to each well and the plate incubated at 4 °C on a platform rocker with occasional agitation. After 20 min, lysates were transferred to chilled microcentrifuge tubes and Laemmli sample buffer (6X; #LB0100, Morganville Scientific, USA), supplemented with β-mercaptoethanol, was added to generate a 1X concentration of Laemmli sample buffer in cell lysate. Lysates were immediately heated to 70 °C for 5 min and subsequently fractionated by SDS-PAGE using 12% polyacrylamide gels, transferred to PVDF membranes, and membranes were blocked with 5% non-fat milk in TBS-T. Membranes were then incubated with mouse anti-mKeima antibodies (#M182-3M, MBL International, Japan) at a 1:1000 dilution in TBS-T, followed by anti-mouse IgG, HRP-linked secondary antibodies (#7076, Cell Signaling Technology, USA) at a 1:10,000 dilution in TBS-T. HRP-conjugated secondary antibody binding, immunoblot imaging, and immunoblot densitometry analysis were performed as described above. The relative cellular levels of processed RPS3-Keima for each condition were calculated by dividing the intensity of immuno-specific bands corresponding to processed Keima (25 kDa) by the intensity of immuno-specific bands corresponding to β-actin (as the loading control).

#### Assessment of relative cellular p62 levels

Cell culture medium was removed from cellular monolayers that were incubated at 37°C and under 5% CO_2_ according to the treatment described in each individual experiment (see figure legends). Cellular monolayers were then incubated at 37 °C and under 5% CO_2_ in nutrient-rich medium (McCoy’s 5A + 10% FBS) containing Actinomycin D (4 μM final concentration; #S8964, SelleckChem, Germany), or, in starvation medium (EBSS) containing Actinomycin D (4 μM final concentration). Use of actinomycin D is recommended when utilizing SQSTM1/p62 as an autophagic flux indicator to prevent synthesis of new SQSTM1/p62 in cells under starvation conditions.^32^ Under conditions where autophagic flux was blocked, cells were also incubated in the presence of chloroquine (60 μM final concentration; #P36240, Thermo Fisher Scientific, USA). After 4 h, cell lysates were collected as previously described.

Cellular lysates were fractionated by SDS-PAGE using 12% polyacrylamide gels, transferred to PVDF membranes, and membranes were blocked with 5% non-fat milk in TBS-T. Membranes were then incubated with rabbit anti-SQSTM1/p62 antibodies (#5114, Cell Signaling Technology, USA) at a 1:3000 dilution in TBS-T, followed by anti-rabbit IgG, HRP-linked secondary antibodies (#7074, Cell Signaling Technology, USA) at a 1:10,000 dilution in TBS-T. HRP-conjugated secondary antibody binding, immunoblot imaging, and immunoblot densitometry analysis were performed as described above. The relative cellular SQSTM1/p62 levels for each condition were calculated by dividing the intensity of immuno-specific bands corresponding to SQSTM1/p62 by the intensity of immuno-specific bands corresponding to β-actin (as the loading control). The percent SQSTM1/p62 remaining was calculated by dividing the relative cellular SQSTM1/p62 levels in cells incubated under starved conditions (EBSS) by the relative cellular SQSTM1/p62 levels in cells incubated under nutrient-rich conditions (McCoy’s 5A + 10% FBS).

#### Assessing mTORC1 activity

Cellular lysates were fractionated by SDS-PAGE using 8% polyacrylamide gels, transferred to PVDF membranes, and membranes were blocked with 5% non-fat milk in TBS-T. Membranes were then incubated with rabbit monoclonal anti-p70 S6K antibodies (#34475, Cell Signaling Technology, USA) at a 1:3000 dilution in TBS-T and rabbit polyclonal anti-Phospho-p70 S6K(Thr389) antibodies (#9205, Cell Signaling Technology, USA) at a 1:1000 dilution in TBS-T, followed by anti-rabbit IgG, HRP-linked secondary antibodies (#7074, Cell Signaling Technology, USA) at a 1:10,000 dilution in TBS-T. HRP-conjugated secondary antibody binding, immunoblot imaging, and immunoblot densitometry analysis were performed as described in “Lysate preparation and immunoblotting” and “Immunoblot densitometric analyses”.

#### Assessing ULK1 dependent activation of autophagy

Cellular lysates were fractionated by SDS-PAGE using 8% polyacrylamide gels, transferred to PVDF membranes, and membranes were blocked with 5% non-fat milk in TBS-T. Membranes were then incubated with rabbit monoclonal anti-ULK1 antibodies (#8054, Cell Signaling Technology, USA) at a 1:3000 dilution in TBS-T, rabbit monoclonal anti-Phospho-ULK1 (Ser757) antibodies (#14202, Cell Signaling Technology, USA) at a 1:3000 dilution in TBS-T, followed by anti-rabbit IgG, HRP-linked secondary antibodies (Cell Signaling Technology) at a 1:10,000 dilution in TBS-T. HRP-conjugated secondary antibody binding, immunoblot imaging, and immunoblot densitometry analysis were performed as described in “Lysate preparation and immunoblotting” and “Immunoblot densitometric analyses”.

#### Assessment of Atg8-Family protein levels

Cellular lysates were fractionated by SDS-PAGE using 12% polyacrylamide gels, transferred to PVDF membranes, and membranes were blocked with 5% low-fat milk in TBS-T. Membranes were then incubated with rabbit monoclonal anti-LC3A antibodies (#4599, Cell Signaling Technology, USA), anti-LC3B antibodies (#3868, Cell Signaling Technology, USA), anti-LC3C antibodies (#14736, Cell Signaling Technology, USA), anti-GABARAP antibodies (#13733, Cell Signaling Technology, USA), anti-GABARAPL1 antibodies (#26632, Cell Signaling Technology, USA), or anti-GABARAPL2 antibodies (#14256, Cell Signaling Technology, USA) at a 1:3000 dilution in TBS-T, followed by anti-rabbit IgG, HRP-linked secondary antibodies (Cell Signaling Technology, USA) at a 1:10,000 dilution in TBS-T. HRP-conjugated secondary antibody binding, immunoblot imaging, and immunoblot densitometry analysis were performed as described above.

#### Generation of LC3B Overexpression Vector

RedTrackCMV-LC3B was generated by restriction cloning. Briefly, the coding sequence for LC3B was cloned from the pSELECT-GFP-hLC3 (#psetz-gfplc3, InvivoGen, USA) plasmid using *Pfu*Ultra II Fusion HS DNA polymerase (#600670, Agilent Technologies, USA) and the following primers to add XhoI (using forward primer) and HindIII (using reverse primer) sites: forward primer 5’-GAT ACT CGA GAT GCC GTC GGA GAA GAC CTT-3’ and reverse primer 5’-CTC AAG CTT TTA CAC TGA CAA TTT CAT CCC G-3’.

The PCR product XhoI-LC3B-HindIII and the RedTrackCMV plasmid were digested with XhoI and HindIII. The resulting fragments were purified using a PCR cleanup kit and ligated at a 3:1 ratio of insert:vector. Ligated plasmid was transformed into chemically competent DH5α *Escherichia coli*. Insertion of the LC3B coding sequence was confirmed by sequencing plasmids isolated from kanamycin resistant transformants.

#### Overexpressing LC3B

HCT116 cells were seeded (approximately 0.04-0.05 × 10^6^ cells per well) on 24-well culture plates and incubated at 37 °C and under 5% CO_2_. 24 h after seeding, monolayers of HCT116 cells incubated at 37 °C and under 5% CO_2_ were transfected with RedTrackCMV (vector control; #50957, Addgene) or RedTrackCMV-LC3B (RedTrackCMV containing the gene encoding LC3B) using Lipofectamine 3000 transfection reagent (#L300015, Thermo Fisher Scientific, USA) as described in the manufacturer’s protocol. Briefly, DNA-lipid complex (50 μL, Opti-MEM medium containing DNA (1 μg), Lipofectamine 3000 Reagent (1.5 μL, provided in kit), and P3000 Reagent (1 μL, provided in kit)) was added directly into each well of the 24-well culture plate. 48 h post-transfection, cell monolayers were visually assessed for transfection efficiency using RFP fluorescence as a readout. Using widefield fluorescence microscopy, approximately greater than 95% RFP-positive cells within the cell monolayer was achieved. Cell monolayers were then treated according to the treatment described in each individual experiment (see figure legends) and harvested for analysis.

#### Generating Mouse Intestinal Organoid Culture

Intestinal organoid culture was established from 20 to 25-week-old female C57BL/6 mice by harvesting intestinal epithelial crypts using a protocol published by Stem Cell Technologies for use with their mouse IntestiCult system. Briefly, 20 to 25-week-old C57BL/6 mice were sacrificed according to ethical guidelines, and approximately 20 cm of small intestine was dissected for intestinal crypt isolation. Intestines were submerged in cold PBS, pH 7.4 and intestinal contents were flushed by injecting 1 mL of cold PBS, pH 7.4 into one end of the intestine. Intestines were splayed open by cutting longitudinally along the entire length of the intestine and opened intestines were washed three times with 1 mL of cold PBS, pH 7.4. The washed intestines were then submerged in fresh, cold PBS, pH 7.4. Intestines were then cut into approximately 2 mm segments, which were transferred to a 50 mL conical tube containing 15 mL of cold PBS, pH 7.4.

Intestinal segments were washed until supernatant became clear, approximately 15 – 20 times, by pipetting up and down three times using a pre-wetted 10 mL serological pipette, letting the segments settle to the bottom of the conical tube by gravity, and removing PBS without disturbing the loose pellet of intestinal segments. For each wash, fresh, cold PBS pH 7.4 (15 mL) was added. Once the supernatant became clear, intestinal segments were resuspended in room temperature Gentle Cell Dissociation Reagent (25 mL; #07174, Stem Cell Technologies, Canada) and incubated on a rocking platform. After rocking for 15 min at room temperature, intestinal segments were allowed to settle at the bottom of the tube, by gravity, and supernatant was removed without disturbing the loose pellet of intestinal segments. Intestinal segments were resuspended in cold 0.1% BSA in PBS pH 7.4 (10 mL) by pipetting up and down 3 times with a pre-wet 10 mL serological pipette. Intestinal segments were allowed to settle, by gravity, and supernatant was collected and filtered through a 70 μm cell strainer (#22-363-548, Fisher Scientific, USA) into a 50 mL conical tube. This filtrate was labeled “Fraction 1”. Subsequent fractions 2 – 4 were collected by resuspending intestinal segments in cold 0.1% BSA in PBS pH 7.4 (10 mL), letting intestinal segments settle by gravity, and collecting and filtering supernatant through a 70 μm cell strainer. All fractions were kept on ice. Fractions contain dissociated intestinal epithelial cells and intestinal crypts. Intestinal crypts were pelleted by centrifugation for 5 min at 290 x g at 4 °C. Supernatants containing dissociated intestinal epithelial cells was decanted, and pellets were resuspended in cold 0.1% BSA in PBS pH 7.4 (10 mL) and then transferred to new, 15 mL conical tubes. Intestinal crypts were again pelleted by centrifugation for 3 min at 200 x g at 4 °C. Supernatants were decanted and pellets were resuspended in cold DMEM/F-12 (10 mL; #10-092-CV, Corning, USA) containing 15 mM HEPES. Fractions enriched with intestinal crypts, as determined by enumerating crypts in each suspension with a hemocytometer, were aliquoted to create three suspensions, each containing 500, 1500, or 3000 crypts. Crypts in each of these three suspensions were pelleted by centrifugation for 5 min at 200 x g at 4 °C. Pellets were resuspended in pre-warmed IntestiCult Organoid Growth Medium (150 μL; #06005, Stem Cell Technologies, Canada). Matrigel (150 μL; #354234, Corning, USA) was added to each suspension to generate a 1:1 ratio of pre-warmed medium and Matrigel. Suspensions were then mixed by pipetting up and down 10 times. Drops of each suspension were spotted into the wells of a pre-warmed 24 well plate (50 μL per drop, one drop per well), and plates were incubated at 37 °C under 5% CO_2_. After 10 min, sufficient time for the Matrigel to solidify, IntestiCult Complete Organoid Medium (750 μL) was added and isolated crypts were incubated at 37 °C under 5% CO_2_. Culture medium was changed every 3 – 4 days by aspirating old medium and adding pre-warmed, fresh medium (750 μL). Every 7 – 10 days, organoids were removed from Matrigel and passaged with a split ratio of 1:4 using the protocol published by Stem Cell Technologies for use with their mouse IntestiCult system. All animal work was reviewed and approved by the Illinois Institutional Animal Care and Use Committee (IACUC protocol #15238). All materials are listed in the Key Resources Table.

#### Assessing LC3 levels in Mouse Intestinal Organoids

Mouse Intestinal Organoids were incubated in 24-well cell culture plates at 37 °C under 5% CO_2_ in IntestiCult Complete Organoid Medium (#06005, Stem Cell Technologies, Canada) containing CHIR-99021 (3 μM final concentration; #72052, Stem Cell Technologies, Canada) and Valproic Acid (VPA, 10 μM final concentration; #72292, Stem Cell Technologies, Canada) to enrich organoids for stem cells.^82^ After approximately 5 – 7 days, stem cell-enriched organoids were further incubated in enrichment medium (IntestiCult Complete Organoid Medium containing CHIR99021 and VPA) in the absence or presence of *Cj*-CDT (10 nM, 5% PBS as carrier control). After 48 h, organoids were removed from Matrigel by incubating at 4 °C in Cell Recovery Solution (750 μL; #354253, Corning, USA) on a platform rocker. After 60 min, Matrigel had dissolved and organoids were pelleted from the suspension by centrifugation for 5 min at 500 x g at 4 °C. Cell pellets were lysed and lysates were prepared and analyzed as described above in “Lysate preparation and immunoblotting,” using anti-α-tubulin antibodies (#2144, Cell Signaling Technology, USA) at a 1:10,000 dilution in TBS-T, followed by anti-Rabbit horseradish peroxidase (HRP)-conjugated secondary antibodies (#7074, Cell Signaling Technology, USA) at a 1:10,000 dilution in TBS-T. For detection of LC3A/B, membranes were incubated with rabbit monoclonal anti-LC3A/B antibodies (#12741, Cell Signaling Technology, USA) at a 1:2000 dilution in TBS-T, followed by anti-rabbit biotinylated secondary antibodies (#14708, Cell Signaling Technology) at a 1:10,000 dilution in TBS-T, followed by anti-biotin horseradish peroxidase (HRP)-conjugated tertiary antibodies (#7074, Cell Signaling Technology, USA), at a dilution of 1:10,000 in TBS-T. HRP-conjugated secondary antibody binding, immunoblot imaging, and immunoblot densitometry analysis were performed as described above.

#### Production of catalytically inactive *Cj*-CDT

The capacity to degrade DNA *in vitro*, as well as induce the DNA damage response, is ablated when a single active site histidine residue (His-157) of *Cj*-CDT, identified by sequence homology to DNase I, was altered by site-directed mutagenesis^42^.

Substitution of the (His-157) residue within the *Cj*-CdtB gene to glycine was performed using site-directed mutagenesis PCR. Briefly, the H157G substitution was generated within the *Cj*-CdtB gene using a pET-15b protein expression vector containing *Cj-CdtB* as a template, *Pfu*Ultra II Fusion HS DNA polymerase (#600670, Agilent Technologies, USA), and primers targeting the *Cj-CdtB* His-157 residue (forward primer: GAT GCT TTT TTC AAT ATC <u>GGT</u> GCT TTA GCT AAT GG, reverse primer: CCA TTA GCT AAA GC<u>A CC</u>G ATA TTG AAA AAA GCA TC). Underlined codon is the location of the His → Gly substitution. Recombinant *Cj*-CDT holotoxin containing catalytically inactive *Cj*-CdtB subunit was expressed and purified exactly as described above for the production of recombinant *Cj*-CDT holotoxin containing catalytically active *Cj*-CdtB subunit.

#### Assessing nuclear localization of *Cj*-CdtB H157G

Cells were seeded, approximately 20,000 cells per well of an 8-well cell culture microscope slide, and incubated at 37 °C and under 5% CO_2_. After 24-36 h, the monolayers were incubated at 37 °C and under 5% CO_2_ in the presence of *Cj*-CDT holotoxin containing catalytically active *Cj*-CdtB or the catalytically inactive *Cj*-CdtB H157G. After 8 h, the cell monolayers were washed three times with PBS pH 7.4. The cell monolayers were then fixed with 4% formaldehyde in PBS pH 7.4. After 15 min at room temperature, fixative was removed, and cell monolayers washed three times with PBS pH 7.4. After washing, cells were permeabilized with 0.1% Triton X-100 in PBS pH 7.4 at room temperature. After 15 min, cell monolayers were then blocked with blocking buffer (5% final concentration of bovine serum albumin in PBS pH 7.4). After 1 h, cell monolayers were washed three times with PBS-T (PBS pH 7.4 containing 0.1% final concentration of Tween-20), 5 min per wash. After washing, cell monolayers were incubated at 4 °C with rabbit polyclonal anti-*Cj*-CdtB antibodies (Yenzym Antibodies, LLC) at a 1:5000 dilution in PBS-T. *Cj*-CdtB antibodies were generated from the immunization of rabbits with a synthesized peptide (CdtB: CDFNRDPSTITSTVDRELANR-amide) by Yenzym Antibodies, LLC (South San Francisco, CA, USA). In order to determine the specificity of *Cj*-CdtB antibodies, immunoblot analysis of purified holotoxin containing *Cj*-CdtA, *Cj*-CdtB, and *Cj*-CdtC subunits using *Cj*-CdtB antibodies was performed and detection of only the *Cj*-CdtB subunit was confirmed. After overnight primary antibody incubation, cell monolayers were washed three times with PBS-T, 5 min per wash. Cell monolayers were then incubated in the presence of secondary antibody (goat anti-rabbit IgG (H+L) Alexa Fluor 488 conjugate; at a 1:1000 dilution in PBS-T; #A11008, Thermo Fisher Scientific, USA). After 1 h at room temperature, cell monolayers were washed three times with PBS-T, 5 min per wash. DAPI (500 ng/mL; #D9542, Sigma-Aldrich, USA) staining solution was then added to the cellular monolayers. After 30 min at room temperature, cell monolayers were washed three times with PBS-T, 5 min per wash. Prolong Gold antifade reagent (#P36930, Thermo Fisher Scientific, USA) was then added to stained monolayers and coverslip applied and sealed with clear nail polish.

#### Assessing alternative agents of DNA damage

HCT116 cells were seeded (approximately 0.04-0.05 × 10^6^ cells per well) on 24-well culture plates and incubated at 37 °C and under 5% CO_2_. 24-36 h after seeding, monolayers of HCT116 cells were incubated at 37 °C and under 5% CO_2_ in the presence of Bleocin (0.1, 1, 10, 100 μg/mL, 5% H_2_O as the carrier control; #203408, Calbiochem, USA), Etoposide (5 μg/mL, 0.1% DMSO as the carrier control; #341205, Calbiochem, USA), or 5-fluorouracil (0.1 and 10 μg/mL, 0.1% DMSO as the carrier control; #sc-29060, Santa Cruz Biotechnology, USA) for up to 24 h.

#### Assessing DNA damage caused by UV-C irradiation

HCT116 cells were seeded (approximately 0.04 – 0.05 × 10^6^ cells per well) on 24-well culture plates and incubated at 37 °C and under 5% CO_2_. 48 h after seeding, HCT116 monolayers, with culture medium removed, were incubated at 37 °C in the absence, or, presence of UV-C radiation in a Hoefer UV-C 500 Crosslinker (254 nm), set to a fluency (defined as power per unit area) of 100 μW/cm^2^ for 5, 25, or 50 s. The dosage of UV-C radiation is calculated as a function of fluency (100 μW/cm^2^) multiplied by the time exposed (5, 25, or 50 s), giving dosages of 5, 25, or 50 J/m^2^, respectively, for monolayers exposed to UV-C radiation. Immediately following exposure to UV-C, fresh medium (McCoy’s 5A + 10% FBS) with Penicillin-Streptomycin (100 units/mL final concentration of Penicillin, 100 μg/mL final concentration of Streptomycin) was added to monolayers and cells were incubated at 37 °C and under 5% CO_2_. After 24 h, monolayers were further incubated in the presence of vehicle control (DMSO, 0.1% final concentration) for LC3-I analysis, or the autophagy inducer CCCP (25 μM) for LC3-II analysis. After an additional 3 h, monolayers were harvested and analyzed for DNA damage response initiation (γ-H2AX), p-p53 (S15), LC3-I, and LC3-II levels as described above.

#### Assessing cell death in DNA damaged cells

HCT116 cells were seeded (approximately 0.01-0.02 × 10^6^ cells per well) on clear bottom black well 96-well culture plates (#3904, Corning, USA) and incubated at 37 °C and under 5% CO_2_. 24-36 h after seeding, monolayers of HCT116 cells were incubated at 37 °C and under 5% CO_2_ in the absence (0.5% DMSO and 10% PBS, as the carrier control) or presence of CDT (1, 10, 100, and 1000 nM), Bleocin (1, 10, 100, and 1000 μg/mL; #203408, Calbiochem, USA), Etoposide (0.5, 5, 50, and 500 μg/mL; #341205, Calbiochem, USA), 5-fluorouracil (0.1, 1, 10, and 100 μg/mL; #sc-29060, Santa Cruz Biotechnology, USA), or staurosporine (1 μM; #ALX-380-014-C100, Enzo Life Sciences, USA). To ensure accurate subtraction of background fluorescence from phenol red in the cell culture medium, negative control medium (0.5% DMSO, 10% PBS) was added to empty wells that contained no cell monolayers. After 24 h, phase-contrast images of monolayers were collected with an OMAX camera (A3550U) using a Zeiss Invertoskop 40C microscope and a Zeiss LD A-Plan 20X phase-contrast objective (1006-591). Immediately after collecting images, PBS (pH 7.4) containing 2X ethidium homodimer-1 (4 μM; #L3224B, Thermo Fisher Scientific, USA) was added to the existing medium in each well at a 1:1 ratio, to yield a final concentration of 2 μM ethidium homodimer-1, and incubated at room temperature protected from light. After 60 min, ethidium homodimer-1 fluorescence was measured using a Synergy 2 microplate reader (#7131000, BioTek) and Gen5 software (BioTek, version 1.11.5) with a 540/25 excitation filter and a 620/40 emission filter. For each biologically independent replicate, background fluorescence (the fluorescence measured in cell-free wells that contained cell culture medium and ethidium homodimer-1) was subtracted from every condition.

#### Inhibiting ATM kinase activity

HCT116 cells were seeded (approximately 0.04-0.05 × 10^6^ cells per well) on 24-well culture plates and incubated at 37 °C and under 5% CO_2_. 24-36 h after seeding, monolayers of HCT116 cells were incubated at 37 °C and under 5% CO_2_ in the absence (DMSO, 0.1% final concentration in cell culture medium as the carrier control) or presence of the ATM inhibitor, KU-55933 (10 μM; #118500, Sigma-Aldrich, USA). After a 1 h pre-treatment with KU-55933, monolayers were further incubated in the absence (DMSO, 0.1% final concentration in cell culture medium) or presence of KU-55933 (10 μM) and in the presence of indicated treatment (see figure legends) for the duration of the experiment, up to 24 h. Inhibition of ATM kinase activity was assessed by comparing levels of phosphorylated p53 at Ser15 (phospho-p53 (Ser15), a target of ATM kinase) in monolayers exposed to *Cj*-CDT in the absence of KU-55933 to the levels of phospho-p53 (Ser15) in monolayers exposed to *Cj*-CDT in the presence of KU-55933 (10 μM) using immunoblot analysis with primary antibodies specific to phospho-p53 (Ser15).

#### Assessing p53 activation and stabilization

Cellular lysates were fractionated by SDS-PAGE using a 12% polyacrylamide gel, transferred to PVDF membranes and membranes were blocked with 5% non-fat milk in TBS-T. Membranes were then incubated with rabbit anti-phospho-p53 (Ser15) antibodies (#9284, Cell Signaling Technology, USA) at a 1:3000 dilution in TBS-T, followed by anti-rabbit IgG, HRP-linked secondary antibodies (#7074, Cell Signaling Technology, USA) at a 1:10,000 dilution in TBS-T. HRP-conjugated secondary antibody binding, immunoblot imaging, and immunoblot densitometry analysis were performed as described above.

#### DNA damage-independent stabilization of p53

HCT116 cells were seeded (approximately 0.04-0.05 × 10^6^ cells per well) on 24-well culture plates and incubated at 37 °C and under 5% CO_2_. 24-36 h after seeding, monolayers of HCT116 cells were incubated at 37 °C and under 5% CO_2_ in the absence (DMSO, 0.1% final concentration in cell culture medium as the carrier control) or presence of the MDM2 inhibitor, Nutlin-3 (20 μM final concentration; #ALX-430-128-M001, Enzo Life Sciences, USA). After 24 h, cell monolayers were then treated according to the treatment described in each individual experiment (see figure legends) and harvested for analysis. Stabilization of p53 as a consequence of Nutlin-3 treatment was assessed through immunoblot analysis for an increase in the total cellular levels of total p53 in monolayers incubated in the presence of Nutlin-3 (20 μM) using primary antibodies specific to p53 (#sc-126, Santa Cruz Biotechnology, USA).

#### Assessing LC3B transcripts

Cell monolayers of HCT116 cells at 37 °C and under 5% CO_2_ were incubated in the absence (PBS pH 7.4 as carrier) or presence of CDT (*Cj*-CDT, 10 nM). After 24 h, total RNA was harvested using the Qiagen RNeasy kit (#74104, Qiagen, Germany) and reverse transcribed into cDNA using SuperScript III Reverse Transcriptase (#18080051, Thermo Fisher Scientific, USA). SYBR green based quantitative RT-PCR was conducted on an Eppendorf Mastercycler realplex^2^ using sense and anti-sense primers to *MAP1LC3B* (coding for human LC3B, F-ACC ATG CCG TCG GAG AAG, R- ATC GTT CTA TTA TCA CCG GGA TTT T),^83^ *CDKN1A* (coding for human p21, F- AGG CAC CGA GGC ACT CAG AG, R- AGT GGT AGA AAT CTG TCA TGC TG),^84^ and *ACTB* (coding for human β-actin, F- CAT GTA CGT TGC TAT CCA GGC, R- CTC CTT AAT GTC ACG CAC GAT) as the housekeeping gene. Data were acquired and analyzed using realplex 2.2 software (Eppendorf).

#### Assessing cellular protein synthesis levels

Cell monolayers of HCT116 were seeded on 12 well plates and incubated at 37 °C with 5% CO_2_. 24-36 h after seeding, monolayers of HCT116 cells were incubated in the absence or presence of CDT (10 nM), or in the presence of puromycin (10 μg/mL) in separate wells as a protein synthesis inhibitor control. After 22 h in the presence of CDT, or 30 min in the presence of puromycin, the medium was removed, and the cells further incubated at 37 °C and under 5% CO_2_ in L-homopropargylglycine (HPG; #1067, Click Chemistry Tools, USA) labeling medium (DMEM-high glucose, no glutamine, no methionine, no cystine, supplemented with 4 mM glutamine, 0.2 mM cystine, and 50 μM HPG) in the continued presence of either CDT or puromycin. After 30 min, the cells were washed with PBS pH 7.4 and trypsinized for 5 min at 37 °C. After confirming detachment of the cells, the enzyme activity of trypsin was neutralized on ice by adding DMEM supplemented with 10% FBS (2 times the volume of trypsin). The cells were resuspended by gentle pipetting and fixed at room temperature by adding the equivalent volume of 8% formaldehyde in PBS pH 7.4, resulting in a 4% final formaldehyde concentration. After 15 min of formaldehyde fixation, the cells were pelleted by centrifugation at 800 *× g* for 1 min and then resuspended at room temperature in 100 μL of a saponin-based permeabilization buffer (1X in 1% BSA in PBS pH 7.4, #C10632, Thermo Fisher Scientific, USA) by gentle vortexing. After 15 min, the HPG molecules incorporated within the cells were fluorescently labeled, at room temperature and protected from light, by adding 100 μl of a freshly prepared fluorescent labeling mixture (1.5 μM FAM Picolyl Azide (#1180, Click Chemistry Tools, USA), 2% Cu-THPTA ligand (#H4050, Lumiprobe, USA), and 10 μM Sodium L-ascorbate (#11140, Sigma-Aldrich, USA) in PBS) and were gently mixed by inverting tubes several times. After 30 min, the cells were pelleted by centrifuging at 800 *× g* for 1 min, and then resuspended in 1% BSA in PBS pH 7.4 at the desired volume for subsequent flow cytometry analyses (LSR II analyzer, BD). 10,000 cells were measured for each analysis. Flow cytometry data were processed using FCS Express software (De Novo Software) and statistics performed using GraphPad Prism 7 (GraphPad Software).

#### Inhibiting proteasome activity

HCT116 cells were seeded (approximately 0.04-0.05 × 10^6^ cells per well) on 24-well culture plates and incubated at 37 °C and under 5% CO_2_. 24-36 h after seeding, monolayers of HCT116 cells were incubated at 37 °C and under 5% CO_2_ in the absence (DMSO, 0.1% final concentration in cell culture medium as the carrier control) or presence of the proteasome inhibitor, MG132 (380 nM final concentration; #M7449, Sigma-Aldrich, USA) and in the presence of the indicated treatment (see figure legends) for the duration of the experiment, up to 24 h. Inhibition of proteasome activity in the presence of MG132 (380 nM) was confirmed through immunoblot analysis for an accumulation of ubiquitinated proteins, as a result of proteasome inhibition, using primary monoclonal antibodies specific to ubiquitin.

#### Immunoprecipitation of Myc-tagged LC3

HCT116 cells were seeded (approximately 0.3 x 10^6^ cells per well) on 6-well cell culture plates and incubated at 37 °C and under 5% CO_2_. After 12 h, cells were transfected with a Myc-tagged LC3 ectopic expression plasmid (pCMV-myc-LC3; #24619, Addgene) using Lipofectamine 3000 transfection reagent (#L300015, Thermo Fisher Scientific, USA) as described in the manufacturer’s protocol. Briefly, 250 μL of DNA-lipid complex (Opti-MEM medium containing 2.5 μg of DNA, 7.5 μL of Lipofectamine 3000 Reagent, and 5 μL of P3000 reagent) were added directly into each well containing HCT116 monolayers, and cells were incubated at 37 °C and under 5% CO_2_. After overnight transfection, cell monolayers were further incubated in the absence or presence of CDT (*Cj-*CDT, 10 nM, 5% PBS as carrier control), and, in the absence or presence of MG132 (380 nM) to prevent degradation of ubiquitinated LC3 by the proteasome.^85,86^ After 12 h, monolayers were further incubated in the absence or presence of TAK-243 (100 nM final concentration; #S8341, SelleckChem, Germany) to block ubiquitination of LC3 by inhibiting ubiquitin activating enzyme activity. After an additional 12 h, monolayers were chilled on ice and then washed twice with ice-cold PBS pH 7.4. To lyse the cells, monolayers were incubated at 4 °C with agitation on a platform rocker in RIPA buffer supplemented with HALT protease and phosphatase inhibitor cocktail, and N-ethylmaleimide (10 mM final concentration; #E1271, Sigma-Aldrich, USA). After 20 min, lysates were transferred to chilled microcentrifuge tubes, and passed through a 25-gauge needle 10 times to homogenize the sample. In order to analyze immunoprecipitation input for each condition, after syringing, a 10% aliquot of whole cell lysate was removed for immunoprecipitation input analysis. To immunoprecipitate Myc-tagged LC3 from lysates, half of each lysate was transferred to a chilled microcentrifuge tube, and mouse anti-Myc antibodies (1:1000 dilution; #2276, Cell Signaling Technology, USA) were added to each tube. For the beads-only control half of each lysate was transferred to a new chilled microcentrifuge tube and no antibodies were added. Lysates were incubated with or without immunoprecipitation antibodies at 4 °C with end-over-end rotation. After a 24 h, 20 μL (approximately 5% of total volume of immunoprecipitation) of pre-washed Protein A magnetic Dynabeads (Invitrogen) were added to each lysate sample, and reactions were incubated at 4 °C with end-over-end rotation. After an additional 24 h, beads were washed three times with ice-cold RIPA buffer, and magnetic bead-associated proteins were eluted from beads by boiling bead suspensions in a denaturing and reducing 1X SDS buffer (approximately 1.5% SDS, 63 mM Tris-HCl, 8% Glycerol, supplemented with β-mercaptoethanol) for 10 minutes.

Elutions were either analyzed via immunoblot analysis immediately, or stored at −80 °C prior to immunoblot analysis in order to preserve ubiquitination of immunoprecipitated proteins. For analysis of high-molecular weight, polyubiquitinated Myc-tagged LC3, elutions from immunoprecipitation (8 μL) were fractionated by SDS-PAGE, using 6% polyacrylamide gels, at 110 V for 90 min, and transferred to PVDF membranes at 30 V for 12 h. For analysis of standard molecular weight Myc-tagged LC3, elutions from immunoprecipitation (8 μL) were fractionated by SDS-PAGE, using 6 % polyacrylamide gels, at 110 V for 90 min, and transferred to PVDF membranes at 30 V for 12 h. For analysis of immunoprecipitation input, Laemmli sample buffer, supplemented with β-mercaptoethanol, was added to generate a 1X concentration of Laemmli sample buffer in input lysates, and input lysates were boiled at 100 °C for 10 min. Input lysates were then fractionated by SDS-PAGE, using 12% polyacrylamide gels, at 110V for 90 min, and transferred to PVDF membranes at 100 V for 90 min. Membranes were blocked with 5% BSA in TBS-T. Membranes were then incubated with rabbit monoclonal anti-Myc antibodies (#2278, Cell Signaling Technology, USA) at a 1:2000 dilution in TBS-T, or rabbit monoclonal anti-Ubiquitin antibodies (#431224, Cell Signaling Technology, USA) at a 1:3000 dilution in TBS-T, followed by anti-rabbit IgG HRP-linked secondary antibodies (#7074, Cell Signaling Technology, USA) at a 1:10,000 dilution in TBS-T. HRP-conjugated secondary antibody binding, immunoblot imaging, and immunoblot densitometry analysis were performed as described above.

#### Immunoprecipitation of endogenous LC3B

HCT116 cells were seeded (approximately 0.3 x 10^6^ cells per well) on 6-well cell culture plates and incubated at 37 °C and under 5% CO_2_. After 24 – 36 h, cell monolayers were further incubated in the absence or presence of CDT (*Cj-*CDT, 10 nM, 5% PBS as carrier control), and, in the absence or presence of MG132 (380 nM) to prevent degradation of ubiquitinated LC3 by the proteasome.^85,86^ After 24 h, monolayers were chilled on ice and then washed twice with ice-cold PBS pH 7.4. After washing the monolayers, RIPA buffer (#89901, Thermo Fisher Scientific, USA) supplemented with HALT protease and phosphatase inhibitor cocktail (#78443, Thermo Fisher Scientific, USA) and N-ethylmaleimide (10 mM final concentration; #E1271, Sigma-Aldrich, USA) was added to each well and the plate was incubated at 4 °C on a platform rocker with occasional agitation. After 20 min, lysates were transferred to chilled microcentrifuge tubes. Lysates, on ice, were then passed through a 25-gauge needle 10 times to fully homogenize the sample. The soluble and insoluble fractions were separated by centrifugation at 14,000 x *g* for 10 min at 4 °C. After centrifugation, soluble fractions were transferred to new tubes. In order to analyze immunoprecipitation input for each condition, an aliquot (approximately 10% of total lysate volume) of lysate soluble fraction was removed for input analysis. To immunoprecipitate endogenous LC3B from lysates, half of each lysate was transferred to a chilled microcentrifuge tube, and mouse anti-LC3B antibodies (#83506, Cell Signaling Technology, USA) were added to each tube. For beads-only control, half of each lysate was transferred to a new chilled microcentrifuge tube and no antibodies were added. Lysates were incubated with and without immunoprecipitation antibodies at 4 °C with end-over-end rotation. After 24 h, 20 μL (approximately 5% of total immunoprecipitation volume) of pre-washed Protein A magnetic Dynabeads (Invitrogen) were added to each lysate sample, and reactions were incubated at 4 °C with end-over-end rotation. After an additional 24 h, beads were washed three times with ice-cold Tris Lysis buffer, and magnetic bead-associated proteins were eluted from beads by boiling bead suspensions in a denaturing and reducing 1X SDS buffer (approximately 1.5% SDS, 63 mM Tris-HCl, 8% Glycerol supplemented with β-mercaptoethanol) for 10 min. Elutions were either analyzed via immunoblot analysis immediately, or stored at −80 °C prior to immunoblot analysis in order to preserve ubiquitination of immunoprecipitated proteins. For analysis of high-molecular weight, polyubiquitinated endogenous LC3B, elutions from immunoprecipitation (8 μL) were fractionated by SDS-PAGE, using 6% polyacrylamide gels, at 110 V for 90 min, and transferred to PVDF membranes at 30 V for 12 h. For analysis of standard molecular weight endogenous LC3B, phagophore-associated SQSTM1/p62, and β-actin, elutions from immunoprecipitation (8 μL) were fractionated by SDS-PAGE, using 12% polyacrylamide gels, at 110 V for 90 min, transferred to PVDF membranes at 100 V for 90 min, blocked with 5% non-fat milk, incubated with rabbit polyclonal anti-SQSTM1/p62 antibodies (1:1000 dilution in TBST; #5114, Cell Signaling Technology, USA), rabbit monoclonal anti-LC3A/B antibodies (1:3000 dilution in TBST; #12741, Cell Signaling Technology, USA), or β-actin (1:10000 dilution in TBST; #4970, Cell Signaling Technology, USA), followed by HRP-linked protein A (1:10000 dilution in TBST; #12291, Cell Signaling Technology, USA). HRP-linked protein A is useful for preventing immunoprecipitation artifacts, such as smears and dense bands, that are often a result of an interaction between primary and secondary antibodies interacting with denatured protein A present in immunoprecipitates. For analysis of immunoprecipitation input, 6X Laemmli sample buffer, supplemented with β-mercaptoethanol, was added to generate a 1X concentration of Laemmli sample buffer in input lysates, and input lysates were boiled at 100 °C for 10 min. Input lysates were then fractionated by SDS-PAGE, using 12% polyacrylamide gels, at 110V for 90 min, and transferred to PVDF membranes at 100 V for 90 min. Membranes were blocked with 5% BSA in TBS-T. Membranes were then incubated with rabbit monoclonal anti-ubiquitin antibodies (#43124, Cell Signaling Technology, USA) at a 1:3000 dilution in TBS-T or rabbit monoclonal anti-LC3A/B antibodies (#12741, Cell Signaling Technology, USA) at a 1:3000 dilution in TBS-T, followed by anti-rabbit IgG HRP-linked secondary antibodies (#7074, Cell Signaling Technology, USA) at a 1:10,000 dilution in TBS-T. HRP-conjugated secondary antibody binding, immunoblot imaging, and immunoblot densitometry analysis were performed as described above.

#### LC3B Immunoprecipitation in Nutlin treated cells

HCT116 cells were seeded (approximately 0.3 x 10^6^ cells per well) on 6-well cell culture plates and incubated at 37 °C and under 5% CO_2_. After 24 – 36 h, cell monolayers were further incubated in the absence (0.1% DMSO as carrier control) or presence of Nutlin-3 (20 μM). After 24 h, MG132 (380 nM final concentration) was added to existing cell culture medium to prevent degradation of ubiquitinated LC3 by the proteasome. After 4 h, monolayers were chilled on ice and then washed twice with ice-cold PBS pH 7.4.

After washing the monolayers, RIPA buffer supplemented with HALT protease and phosphatase inhibitor cocktail and N-ethylmaleimide (10 mM final concentration) was added to each well and the plate was incubated at 4 °C on a platform rocker with occasional agitation. After 20 min, lysates were transferred to chilled microcentrifuge tubes. Lysates, on ice, were then passed through a 25-gauge needle 10 times to fully homogenize the sample. Endogenous LC3B immunoprecipitation was performed as described above in “Immunoprecipitation of endogenous LC3B for analysis of ubiquitinated LC3”. For input analysis, a portion of the initial whole cell lysate (10%) was reserved, 6X Laemmli sample buffer, supplemented with β-mercaptoethanol, was added to generate a 1X concentration of Laemmli sample buffer in input lysates, and input lysates were fractionated by SDS-PAGE, using 12% polyacrylamide gels, at 110V for 90 min, and transferred to PVDF membranes at 100 V for 90 min. Membranes were blocked with 5% milk in TBS-T. Total cellular p53, LC3A/B, or β-actin levels were determined by immunoblot analysis using primary antibodies specific to p53 (#sc-126, Santa Cruz Biotechnology, USA), LC3A/B (#12741, Cell Signaling Technology, USA), and β-actin (#4970, Cell Signaling Technology, USA).

#### Inhibiting ATM kinase in *Cj*-CDT pre-treated cells

HCT116 cells were seeded (approximately 0.04-0.05 × 10^6^ cells per well) on 24-well culture plates and incubated at 37 °C and under 5% CO_2_. 24-36 h after seeding, monolayers of HCT116 cells were incubated at 37 °C and under 5% CO_2_ in the absence or presence of *Cj*-CDT (10 nM). After 24 h, culture medium containing toxin was removed and monolayers were further incubated in the absence (DMSO, 0.1% final concentration in cell culture medium as the carrier control) or presence of the ATM inhibitor, KU-55933 (10 μM final concentration). After 4 h, cell lysates were prepared and analyzed as described above in “Lysate preparation and immunoblotting” and “Immunoblot densitometric analysis”.

#### Assessing CtIP and cleaved-caspase 3 levels

HCT116 cells were seeded (approximately 0.04-0.05 × 10^6^ cells per well) on 24-well culture plates and incubated at 37 °C and under 5% CO_2_. 24 h after seeding, monolayers of HCT116 cells incubated at 37 °C and under 5% CO_2_ were transfected with RedTrackCMV (vector control; #50957, Addgene) or RedTrackCMV-LC3B (RedTrackCMV containing the gene encoding LC3B) using Lipofectamine 3000 transfection reagent as described in the manufacturer’s protocol. Briefly, DNA-lipid complex (50 μL, Opti-MEM medium containing DNA (1 μg), Lipofectamine 3000 Reagent (1.5 μL), and P3000 Reagent (1 μL)) was added directly into each well of the 24-well culture plate. Monolayers were observed by live immunofluorescence microscopy to have greater than 95% RFP-positive cells. 24 h post-transfection, cell monolayers were incubated in the absence (5% PBS) or presence of *Cj*-CDT (10 nM, 5% PBS). After 24 h, cell monolayers were harvested for analysis. Cellular lysates were fractionated by SDS-PAGE (12% gels), transferred to polyvinylidene difluoride (PVDF) membranes, and membranes were blocked with 5% non-fat milk in TBS-T. Membranes were then incubated with rabbit monoclonal anti-CtIP antibodies (1:3000 dilution in TBS-T; #9201, Cell Signaling Technology, USA), rabbit monoclonal anti-cleaved caspase-3 (1:3000 dilution in TBS-T; #9661, Cell Signaling Technology, USA), or rabbit monoclonal anti-GAPDH antibodies (1:20,000 dilution in TBS-T; #5174, Cell Signaling Technology, USA), followed by anti-rabbit IgG, HRP-linked secondary antibodies (1:10,000 dilution in TBS-T; #7074, Cell Signaling Technology, USA). HRP-conjugated secondary antibody binding, immunoblot imaging, and immunoblot densitometry analysis were performed as described above under “Immunoblot densitometric analyses”.

#### Assessing apoptosis in cells overexpressing LC3B

HCT116 cells were seeded (approximately 0.05 x 10^6^ cells per well) on 12-well cell culture plates and incubated at 37 °C and under 5% CO_2_. After 24 h, monolayers were transfected with either RedTrackCMV (vector control) or RedTrackCMV-LC3B (RedTrackCMV containing the gene encoding LC3B) using Lipofectamine 3000 transfection reagent, as described in their protocol. Briefly, 100 μL of DNA-lipid complex (Opti-MEM medium containing 2 μg of DNA, 3 μL of Lipofectamine 3000 Reagent, and 2 μL of P3000 reagent) were added directly into each well containing HCT116 monolayers, and cells were incubated at 37 °C and under 5% CO_2_. As a point of comparison, additional monolayers of HCT116 cells were not subjected to transfection. After overnight transfection, monolayers were washed 3X with cell culture medium to remove cells that detached during transfection and monolayers were then further incubated in the absence or presence of CDT (*Cj*-CDT, 10 nM, 5% PBS as carrier control). After 24 h, cell culture medium was removed, and monolayers were washed once with cold 1X PBS pH 7.4. Detached cells were collected by pelleting culture medium and PBS wash in a microcentrifuge at 500 x*g* and 4 °C for 5 min. Adherent cells were collected by incubating at 37 °C in the presence of trypsin. After 7 min, approximately 10 volumes of culture medium containing 10% FBS was added to inactivate trypsin. Adherent cells collected by trypsinization were combined with detached cells (collected before trypsin treatment), and the concentration of cells per mL was determined using a hemocytometer. Cells were stained using Annexin-V PI kit (#556547, BD Pharmingen, USA) as described in the kit manufacturer’s protocol.

Briefly, collected cells were pelleted in a microcentrifuge at 500 x*g* and 4 °C. Cell pellets were washed once with PBS (pH 7.4) to remove calcium chelator EDTA (present in trypsin) which can interfere with Annexin-V binding. After 5 min, the cell pellets were resuspended in 1X Annexin binding buffer to a final concentration of approximately 1.0 x 10^6^ cells/mL (as determined by hemocytometer counting). 100 μL of cell suspensions (approximately 1.0 x 10^5^ cells/mL) were transferred to flow cytometry tubes and combined with 5 μL of Annexin-V stain, and cells were incubated at room temperature (approximately 25 °C) and protected from light. After 20 min, cells were analyzed using a BD FACS Canto II flow cytometry analyzer using a 488 nm excitation laser. Red fluorescence emission (representing transfected cells expressing mRFP1 from RedTrackCMV) was measured to isolate transfected cells, and green fluorescence emission (Annexin-V-FITC) was measured to identify apoptotic cells. Compensation was calculated using (1) non-transfected, unstained cells to determine baseline autofluorescence, (2) transfected, unstained cells (mRFP1 overexpressing cells that were not stained with Annexin-V-FITC) to determine the red fluorescence emission spectra, and (3) non-transfected cells that were treated with Staurosporine (1 μM, 24 h) to induce cell death and stained with Annexin-V-FITC to determine green fluorescence emission spectra. 10,000 individual cells in the mRFP^+^ population were analyzed. Gates were drawn to isolate (1) singlet cells, (2) mRFP^+^ cells, and (3) Annexin-V-FITC^HIGH^ versus Annexin-V-FITC^LOW^ cells. Flow cytometry data were processed using FCS Express software (De Novo Software). Background cell death as a consequence of transfection, as determined by the amount of cell death in the monolayers that were transfected with RedTrackCMV control plasmids that were not incubated in the presence of CDT, was subtracted from the determined percentage values of Annexin-V-FITC^HIGH^ cells to give normalized cell death percentages. Statistics were performed using GraphPad Prism 7 (GraphPad Software).

## QUANTIFICATION AND STATISTICAL ANALYSIS

### Statistical Analyses

Data are presented as mean ± SD (standard deviation). All statistical analysis was performed using GraphPad Prism 7. Analysis of statistical differences between two groups was determined using unpaired, two-tailed Student’s *t*-test. For comparison of three or more groups with one independent variable, analysis of statistical differences was performed using one-way ANOVA followed by Dunnett’s post-hoc test. For comparison of three or more groups with two independent variables, analysis of statistical differences was performed using two-way ANOVA followed by Fisher’s LSD post-hoc test. Figure legends describe the statistical test used for presented data.

Statistical significance was determined at *P* < 0.05, α = 0.05. Choice of sample size, which was at least 3 biological replicates (n=3), ensured adequate power (1-β > 0.8) to detect an effect size of d ≥ 1.5.

