## Supplemental Materials for "Autophagy suppression in DNA damaged cells occurs through a newly identified p53-proteasome-LC3 axis"

1 # SUPPLEMENTAL MATERIALS

#### Figure S1

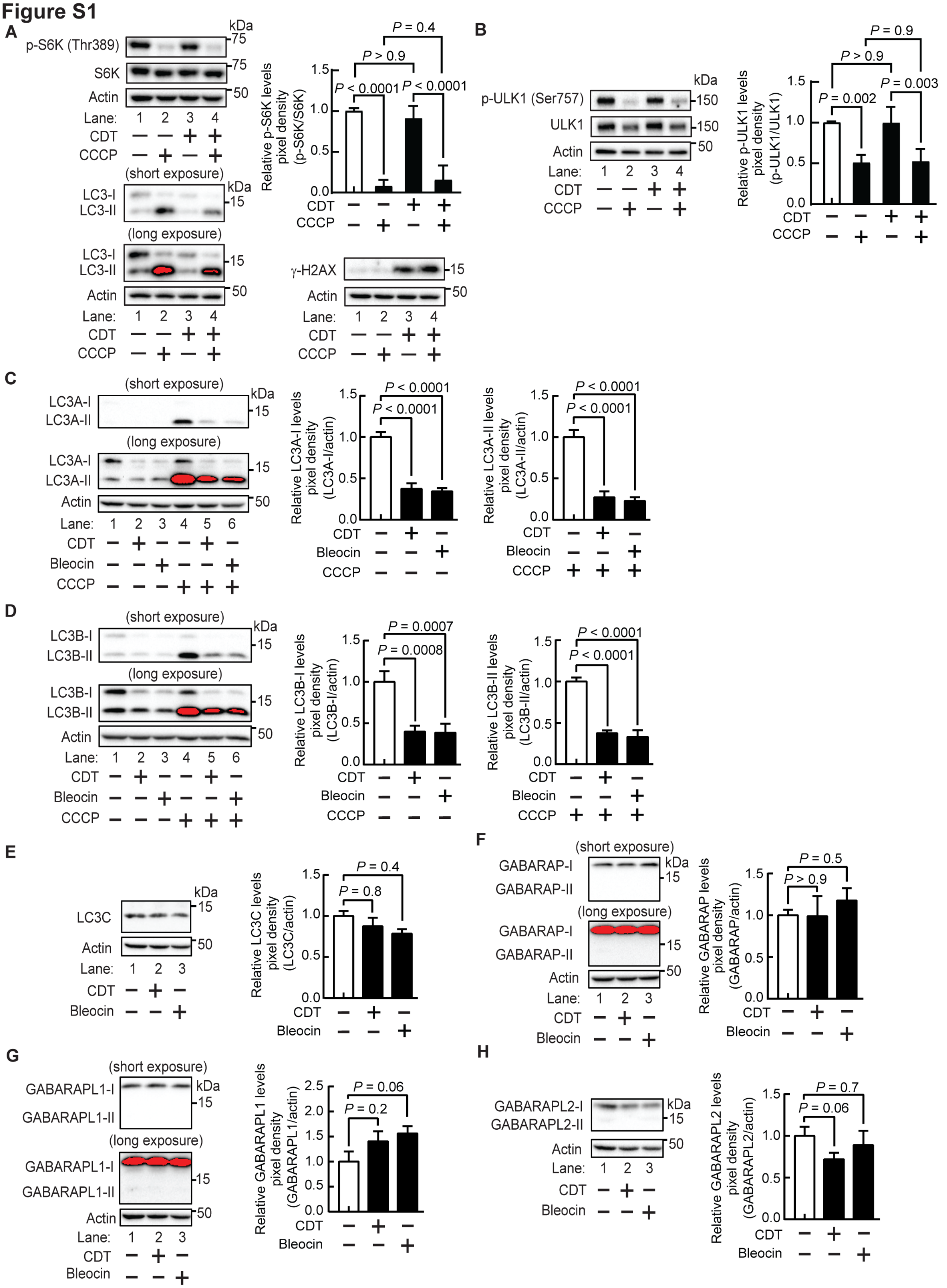


2

**Figure S1. *Cj*-CDT-mediated inhibition of autophagy occurs downstream of mTOR and Ulk1 signaling and is limited to LC3B and LC3A**

Monolayers of HCT116 cells at 37 °C and under 5% $CO_2$ were incubated in the absence or presence of *Cj*-CDT (10 nM) (**A** to **H**) or Bleocin (10 μg/mL) (**C** to **G**). After 24 h, monolayers were further incubated at 37 °C in the absence or presence of the autophagy inducing compound CCCP (25 μM). After an additional 3 h, cell monolayers were lysed, and the lysates analyzed by immunoblot analysis for relative levels of p-S6K (Thr389), total S6K, LC3-I, LC3-II, γ-H2AX (**A**), p-ULK1 (Ser757), total Ulk1 (**B**), LC3A-I, LC3A-II (**C**), LC3B-I, LC3B-II (**D**), LC3C-I, LC3C-II (**E**), GABARAP-I, GABARAP-II (**F**), GABARAPL1-I, GABARAPL1-II (**G**), GABARAPL2-I, GABARA L2-II (**H**), and β-actin (**A** to **H**) (as a loading control). Immunoblots shown are representative of the immunoblots collected from 3 biologically independent experiments (n=3). Densitometric analyses of immunoblots collected from 3 biologically independent experiments (n=3) were combined, and relative cellular levels of p-S6K (Thr389) (**A**) or p-Ulk1 (Ser757) (**B**) were calculated relative to total cellular S6K (**A**) or Ulk1 (**B**) levels, respectively, and relative cellular levels of each Atg8 ortholog were calculated relative to β-actin (**C** to **H**). The data represented by the white bars (monolayers pre-incubated in the absence of *Cj*-CDT) were assigned an arbitrary value of 1.0 (**A** to **H**). Error bars represent standard deviations of data combined from 3 biologically independent experiments (n=3) (**A** to **H**). Statistical analysis of the data was conducted using two-way ANOVA followed by Fisher's LSD post-hoc test (**A** and **B**), or one-way ANOVA, followed by Dunnett's post-hoc test (**C** to **H**). Data are presented as mean ± SD. $P < 0.05$ indicates statistical significance ($\alpha = 0.05$).

**Figure S2**

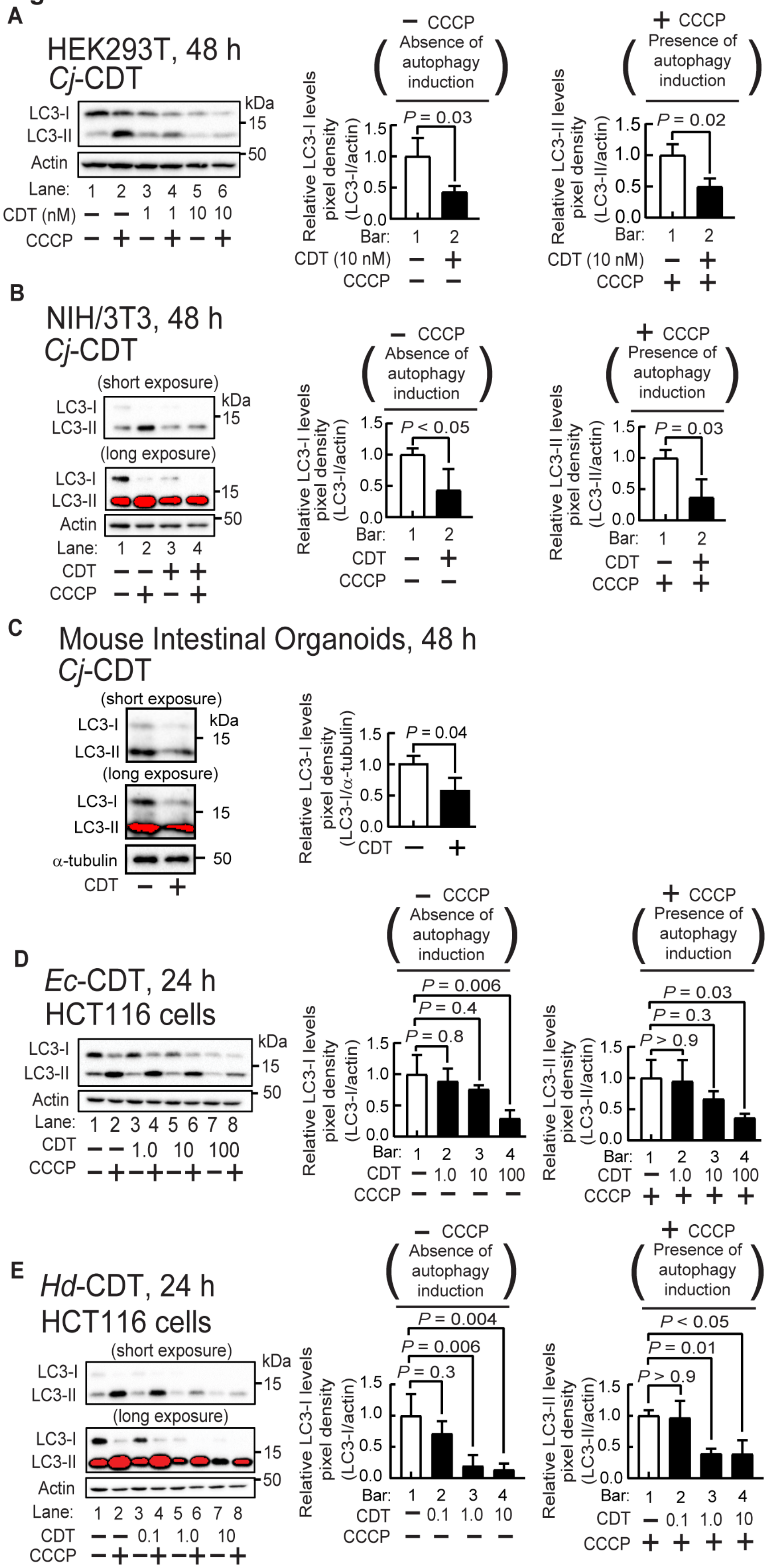
A
HEK293T, 48 h
Cj-CDT
LC3-I
LC3-II
Actin
kDa
15
50
Lane: 1 2 3 4 5 6
CDT (nM) − − 1 1 10 10
CCCP − + − + − +
− CCCP
Absence of autophagy induction
Relative LC3-I levels pixel density (LC3-I/actin)
P = 0.03
Bar: 1 2
CDT (10 nM) − +
CCCP − −
+ CCCP
Presence of autophagy induction
Relative LC3-II levels pixel density (LC3-II/actin)
P = 0.02
Bar: 1 2
CDT (10 nM) − +
CCCP + +
B
NIH/3T3, 48 h
Cj-CDT
(short exposure)
(long exposure)
LC3-I
LC3-II
Actin
kDa
15
50
Lane: 1 2 3 4
CDT − − + +
CCCP − + − +
− CCCP
Absence of autophagy induction
Relative LC3-I levels pixel density (LC3-I/actin)
P < 0.05
Bar: 1 2
CDT − +
CCCP − −
+ CCCP
Presence of autophagy induction
Relative LC3-II levels pixel density (LC3-II/actin)
P = 0.03
Bar: 1 2
CDT − +
CCCP + +
C
Mouse Intestinal Organoids, 48 h
Cj-CDT
(short exposure)
(long exposure)
LC3-I
LC3-II
α-tubulin
kDa
15
50
CDT − +
Relative LC3-I levels pixel density (LC3-I/α-tubulin)
P = 0.04
CDT − +
D
Ec-CDT, 24 h
HCT116 cells
LC3-I
LC3-II
Actin
kDa
15
50
Lane: 1 2 3 4 5 6 7 8
CDT − − 1.0 10 100
CCCP − + − + − + − +
− CCCP
Absence of autophagy induction
Relative LC3-I levels pixel density (LC3-I/actin)
P = 0.006
P = 0.4
P = 0.8
Bar: 1 2 3 4
CDT − 1.0 10 100
CCCP − − − −
+ CCCP
Presence of autophagy induction
Relative LC3-II levels pixel density (LC3-II/actin)
P = 0.03
P = 0.3
P > 0.9
Bar: 1 2 3 4
CDT − 1.0 10 100
CCCP + + + +
E
Hd-CDT, 24 h
HCT116 cells
(short exposure)
(long exposure)
LC3-I
LC3-II
Actin
kDa
15
50
Lane: 1 2 3 4 5 6 7 8
CDT − − 0.1 1.0 10
CCCP − + − + − + − +
− CCCP
Absence of autophagy induction
Relative LC3-I levels pixel density (LC3-I/actin)
P = 0.004
P = 0.006
P = 0.3
Bar: 1 2 3 4
CDT − 0.1 1.0 10
CCCP − − − −
+ CCCP
Presence of autophagy induction
Relative LC3-II levels pixel density (LC3-II/actin)
P < 0.05
P = 0.01
P > 0.9
Bar: 1 2 3 4
CDT − 0.1 1.0 10
CCCP + + + +

26

**Figure S2. CDT-dependent reduction in cellular LC3-I and LC3-II levels is recapitulated in HEK293T cells, NIH/3T3 cells, mouse intestinal organoids, and by CDTs from different bacteria**

Monolayers of HEK293T cells (**A**), NIH/3T3 cells (**B**), murine intestinal organoids (**C**), or HCT116 cells (**D** and **E**), were incubated at 37 °C and under 5% $CO_2$, in the absence (PBS pH 7.4 as the carrier control) or presence of 1 nM *Cj*-CDT (**A**) or 10 nM *Cj*-CDT (**A** to **C**), *Ec*-CDT (1, 10, 100 nM) (**D**), or *Hd*-CDT (0.1, 1, 10 nM) (**E**). After 48 h (**A** to **C**) or 24 h (**D** and **E**), the monolayers were washed, and further incubated with CCCP (25 μM) (**A, B, D,** and **E**), or, lysed (**C**). After 3 h, lysates were analyzed by immunoblot analysis for relative levels of LC3-I, LC3-II, and β-actin (**A, B, D,** and **E**) or α-tubulin (**C**). Immunoblots are representative from 3 biologically independent experiments (n=3). Densitometric analyses from 3 biologically independent experiments were combined and relative cellular LC3-I or LC3-II levels were calculated from dividing the intensity of immuno-specific bands corresponding to LC3-I or LC3-II by the intensity of immuno-specific bands corresponding to β-actin (**A, B, D,** and **E**) or α-tubulin (**C**). The data represented by bar 1 (empty bar) were assigned an arbitrary value of 1.0. Error bars represent standard deviations. Two-tailed, unpaired Student's *t*-tests (**A** to **C**), or one-way Anova, followed by Dunnett's post hoc test (**D** and **E**), were used. Data are presented as mean ± SD. $P < 0.05$ indicates statistical significance ($\alpha = 0.05$).

Figure S3

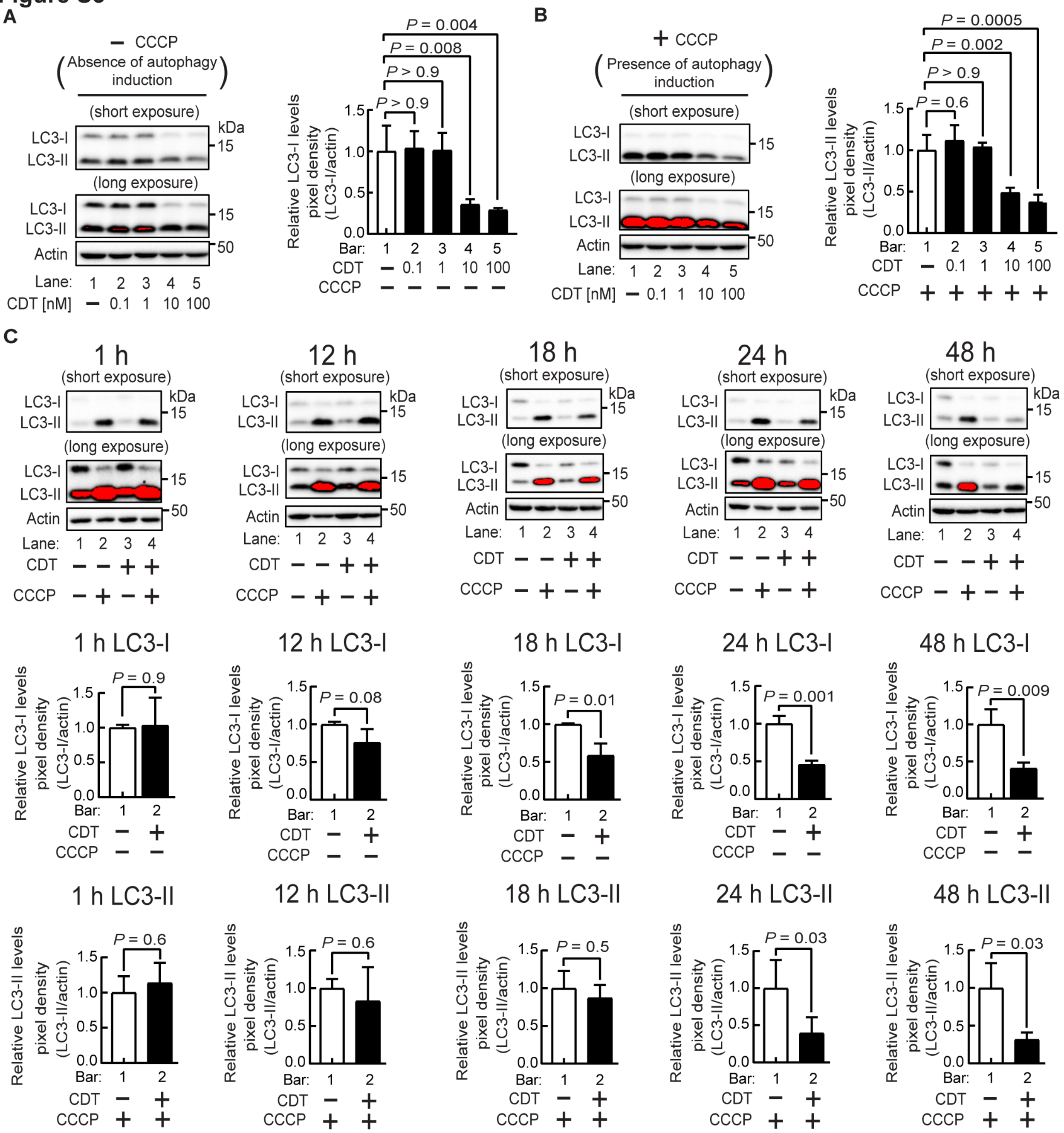


**Figure S3. CDT dose-dependent reduction in cellular LC3-I levels is associated with CDT-dependent impairment of the elevation in LC3-II levels normally observed during autophagy induction, related to Figure 1**

Monolayers of HCT116 cells were incubated in the absence (PBS pH 7.4 as the carrier control) or presence of *Cj*-CDT (0.1, 1.0, 10, and 100 nM) (**A** and **B**), or (10 nM) (**C**). After 24 h (**A** and **B**), or at the indicated timepoints (1, 12, 18, 24, or 48 h) (**C**), cell monolayers were further incubated, in the absence of toxin, and, as indicated, the autophagy inducer CCCP (25 μM). After an additional 3 h, lysates were analyzed by immunoblot analysis for relative levels of LC3-I, LC3-II, and β-actin (**A** to **C**). All immunoblots shown are representative of the immunoblots collected from 3 biologically independent experiments (n=3). The position of the molecular weight (kDa) markers (not shown) are indicated by tick marks to the right of the immunoblots. Densitometric analyses of immunoblots collected from 3 biologically independent experiments were combined and the relative cellular LC3-I or LC3-II levels were calculated by dividing the intensity of immuno-specific bands corresponding to LC3-I or LC3-II by the intensity of immuno-specific bands corresponding to β-actin (as the loading control). The data represented by bar 1 (empty bar) were assigned an arbitrary value of 1.0. Error bars represent standard deviations of data combined from 3 biologically independent experiments (n=3). Statistical analyses of the data were conducted using one-way Anova, followed by Dunnett's post hoc test (**A** and **B**), or two-tailed, unpaired Student's *t*-tests (**C**). Data are presented as mean ± SD. $P < 0.05$ indicates statistical significance ($\alpha = 0.05$).

**Figure S4**

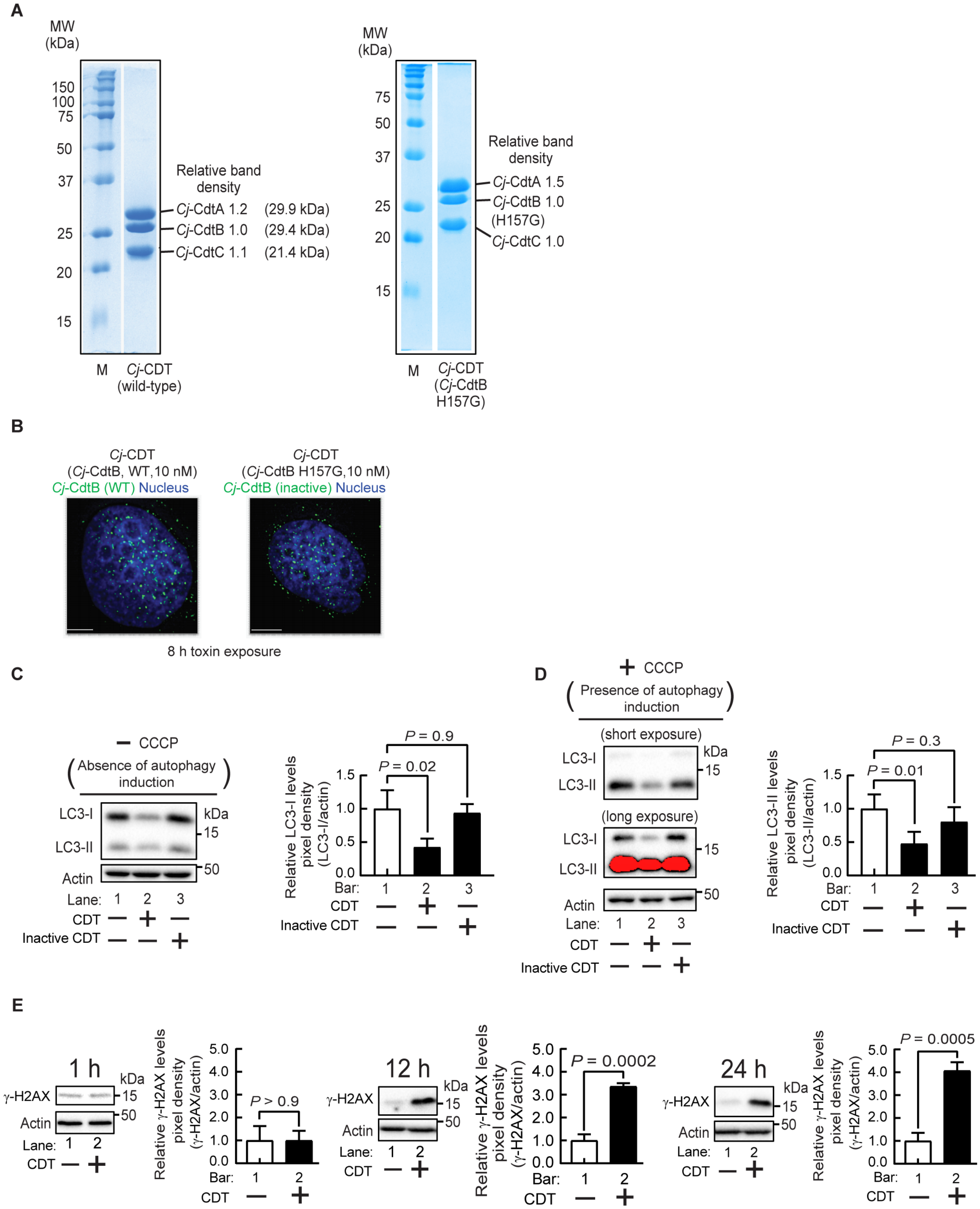


**Figure S4. CDT DNase activity is important for CDT-dependent reduction in both cellular LC3-I and LC3-II levels, related to Figure 1**

SDS-PAGE analysis of wild-type, or, catalytically inactive *Cj*-CDT holotoxin (**A**). Relative band intensities are normalized to the band intensity of the *Cj*-CdtB, or, *Cj*-CdtB-H157G subunit, respectively, which were each assigned an arbitrary value of 1.0 (**A**). HCT116 cells were incubated in the presence of either wild-type CDT (*Cj*-CDT, 10 nM) or catalytically inactive CDT (10 nM) (**B**). After 8 h, cells were fixed and imaged for intracellular *Cj*-CdtB or *Cj*-CdtB-H157G. Images are representative of those collected from three biologically independent experiments (n=3) collected at 40X magnification. White scale bars indicate 2 $\mu$m (**B**). HCT116 cells were incubated for 24 h with *Cj*-CDT (10 nM), or *Cj*-CdtB (H157G) (10 nM), and then further incubated with CCCP (25 $\mu$M) (**C** and **D**). After an additional 3 h, cell lysates were analyzed by immunoblot analysis for relative levels of LC3-II, LC3-I, and $\beta$-actin (**C** and **D**). HCT116 cells were incubated with *Cj*-CDT (10 nM) for 1, 12, or 24 h (**E**). At the indicated timepoints, cell lysates were analyzed by immunoblot analysis for relative levels of $\gamma$-H2AX and $\beta$-actin (**E**). Densitometric analyses were combined from 3 biologically independent experiments (n=3). Data represented by white bars were assigned an arbitrary value of 1.0. Error bars represent standard deviations. Statistical analyses of the data were conducted using one-way Anova, followed by Dunnett's post hoc test (**C** and **D**), or, unpaired Student's *t*-test (**E**). Data are presented as mean ± SD. $P < 0.05$ indicates statistical significance ($\alpha = 0.05$).

Figure S5

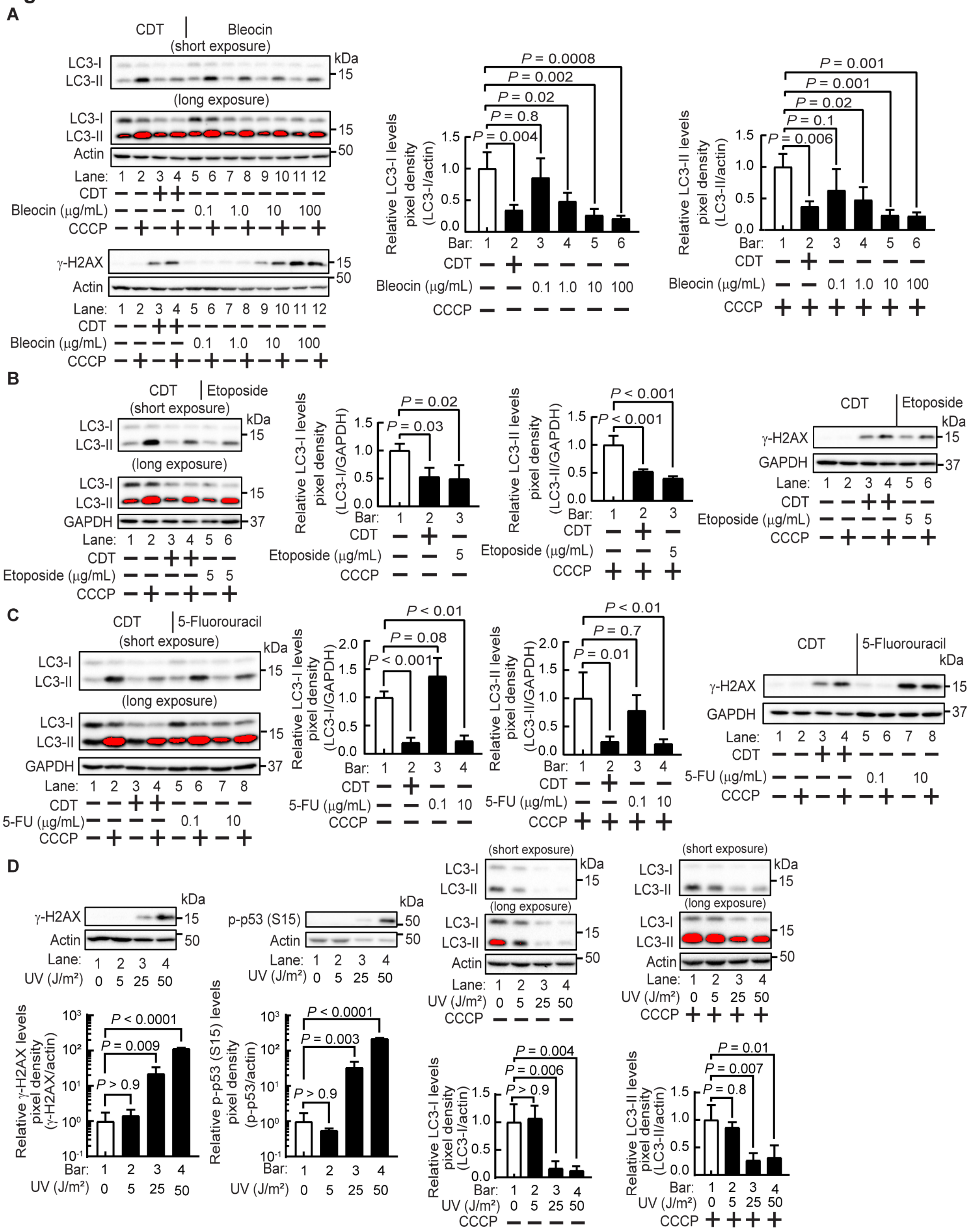


91

92 **Figure S5. Diverse mechanisms of DNA damage recapitulate the *Cj*-CDT-**

93 **dependent reduction in cellular LC3-I and LC3-II levels, related to Figure 1**

HCT116 cells were incubated in the absence or presence of *Cj*-CDT (10 nM) (**A** to **C**), Bleocin (0.1, 1, 10, 100 μg/mL) (**A**), Etoposide (5 μg/mL) (**B**), or 5-Fluorouracil (0.1, 10 μg/mL) (**C**). Alternatively, monolayers were exposed to UV-C radiation at dosages of 5, 25, and 50 $J/m^2$ (**D**). After 24 h, cell monolayers were further incubated in the absence or presence of CCCP (25 μM) (**A** to **D**). After an additional 3 h, cell lysates were analyzed by immunoblot analysis for relative levels of LC3-I, LC3-II, γ–H2AX, β-actin (**A** and **D**), GAPDH (**B** and **C**), and p-p53 (S15) (**D**). Immunoblots shown are representative of the immunoblots collected from 3 biologically independent experiments (n=3). Densitometric analyses of immunoblots collected from 3 biologically independent experiments (n=3) were combined, and relative cellular LC3-I or LC3-II levels were calculated by dividing the intensity of immuno-specific bands corresponding to LC3-I or LC3-II by the intensity of immuno-specific bands corresponding to β-actin or GAPDH. The data represented by the white bars (monolayers pre-incubated in the absence of *Cj*-CDT) were assigned an arbitrary value of 1.0. Error bars represent standard deviations. Statistical analyses of the data were conducted using one-way ANOVA, followed by Dunnett's post-hoc test. Data are presented as mean ± SD. $P < 0.05$ indicates statistical significance ($\alpha = 0.05$).

#### Figure S6

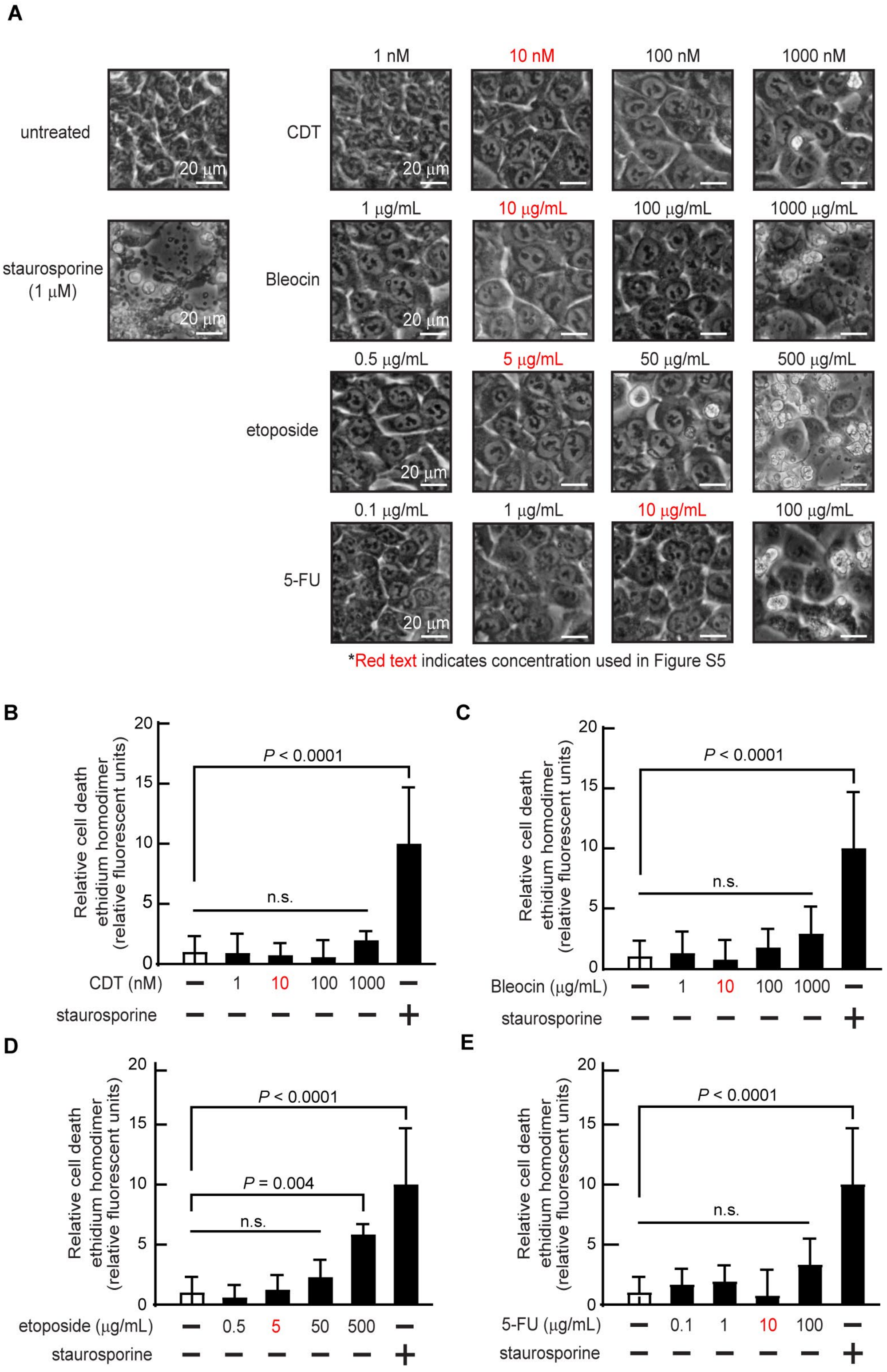


**Figure S6. Cell death is not observed at concentrations of DNA damaging agents associated with LC3 reduction and H2AX activation, related to Figure S5**

Monolayers of HCT116 cells at 37 °C and under 5% $CO_2$ were incubated in the absence (0.5% DMSO, as carrier control) or presence of *Cj*-CDT (1, 10, 100, 1000 nM) (**A** and **B**), Bleocin (1, 10, 100, 1000 $\mu$g/mL) (**A** and **C**), etoposide (0.5, 5, 50, 500 $\mu$g/mL) (**A** and **D**), 5-fluorouracil (5-FU; 0.1, 1, 10, 100 $\mu$g/mL) (**A** and **E**), or staurosporine (1 $\mu$M) (**A** to **E**). After 24 h, phase-contrast microscopy images were collected and relative cell death was evaluated by an ethidium homodimer membrane permeability assay. Microscopy images are representative of those collected from 4 biologically independent experiments (n=4) collected at 20X magnification. White scale bars from representative indicate 20 $\mu$m. Relative fluorescence intensities of ethidium homodimer-1 (EthD-1) incorporation collected from 4 biologically independent experiments (n=4) were combined, and relative cell death was calculated by dividing the EthD-1 fluorescence intensity in treated cells by the EthD-1 fluorescence intensity in untreated cells, which represents basal cell death (**B** to **E**). The data represented by the white bars (monolayers incubated in the absence of treatment) were assigned an arbitrary value of 1.0 (**B** to **E**). The data for the positive control condition (monolayers incubated in the presence of staurosporine) and negative control condition (monolayers incubated in the absence of treatment, white bars) were re-used for each graph (**B** to **E**), as multiple experiments (all 18 conditions) were performed at the same time in a shared 96-well plate for each biologically independent replicate. The data were separated into individual graphs to simplify presentation. Error bars represent standard deviations of data combined from 4 biologically independent experiments (n=4) (**B** to **E**). Statistical analysis of the data was conducted using one-way Anova, followed by Dunnett’s post-hoc test (**B** to **E**). Data are presented as mean ± SD. $P < 0.05$ indicates statistical significance ($\alpha = 0.05$).

**Figure S7**

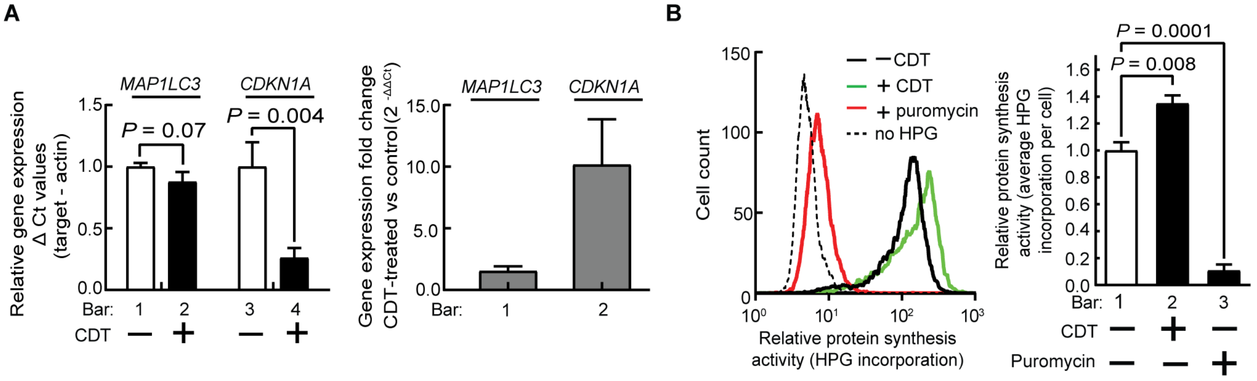


**Figure S7. LC3B mRNA and total cellular protein synthesis are not reduced in cells incubated with CDT, related to Figure 5**

HCT116 cells were incubated in the absence or presence of *Cj*-CDT (10 nM). (**A**) After 24 h, total RNA was analyzed by quantitative RT-PCR analysis for LC3B mRNA and CDKN1A (coding for human p21) as a control for DNA damage-mediated transcriptional regulation. Error bars represent standard deviations from 3 biologically independent experiments (n = 3), with 4 technical replicates per biological experiment. Statistical analyses of the data were conducted using two-tailed, unpaired Student's *t*-tests. $P < 0.05$ indicates statistical significance ($\alpha = 0.05$). (**B**) After 24 h, the cells were further incubated in the L-homopropargylglycine (HPG) labeling medium in the absence or presence of puromycin (10 $\mu$g/mL final concentration). HPG incorporation was determined using flow cytometry. The histograms shown are representative of the histograms collected from 3 biologically independent experiments (n=3). For each of the 3 independent biological replicates, the mean fluorescence intensities of Alexa Fluor 488 labeled cells from 2 technical replicates (each analyzing 10,000 cells) for each treatment were averaged. The data represented by the black bars were rendered relative to the white bar, which was assigned an arbitrary value of 1.0. Error bars represent standard deviations. Statistical analyses of the data were conducted using one-way Anova, followed by Dunnett's post hoc test. Data are presented as mean ± SD. $P < 0.05$ indicates statistical significance ($\alpha = 0.05$).

**Figure S8**

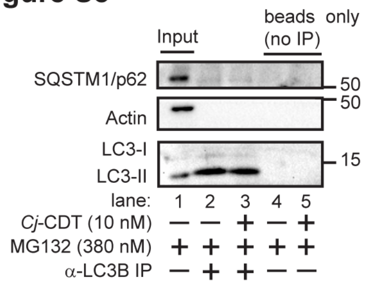


**Figure S8. Phagophore-associated proteins are not detected in LC3-specific immunoprecipitates, related to Figure 5**

HCT116 cells were incubated in the absence or presence of *Cj*-CDT (10 nM), in the presence of MG132 (380 nM). After 24 h, cell lysates were examined for LC3-I, LC3-II, SQSTM1/p62, and β-actin (lane 1). Alternatively, cell lysates were incubated in the absence (“beads only”, lanes 4 and 5) or presence (immunoprecipitation, lanes 2 and 3) of LC3-specific antibodies, in the presence of protein-A-conjugated magnetic beads. Immunoprecipitates were examined for LC3-I, LC3-II, SQSTM1/p62, and β-actin. Immunoblots are representative from data collected from at least 3 biologically independent experiments (n=3).

**Figure S9**

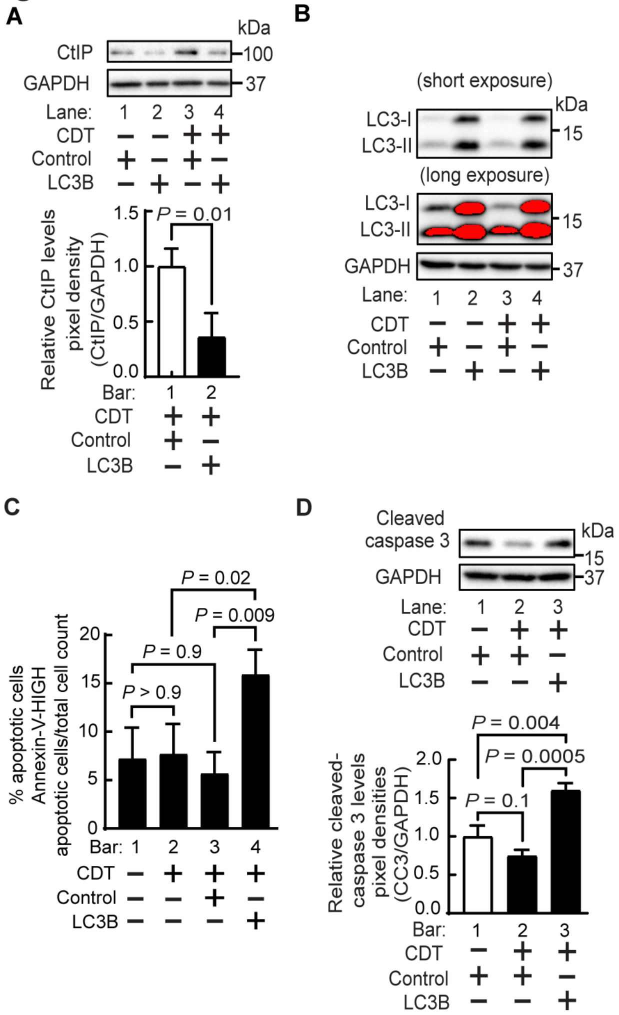


**Figure S9. CDT-mediated autophagy reduction promotes cell viability during DNA damage repair, related to Figure 7**

HCT116 cells transfected with RedTrackCMV (control) or RedTrackCMV-LC3B were incubated in the absence or presence of *Cj*-CDT (**A** to **D**). After 24 h, lysates were analyzed by immunoblot analysis for relative levels of CtIP (**A**), LC3-I, LC3-II (**B**), cleaved caspase 3 (**D**), and GAPDH (**A** and **B**, **D**). Densitometric analyses of immunoblots collected from 3 biologically independent experiments were combined (n = 3). Error bars represent standard deviations. Statistical analyses of the data were conducted using unpaired, two-tailed Student's *t*-test (**A**) or one-way Anova followed by Dunnett's post hoc test (**D**). Data are presented as mean ± SD. $P < 0.05$ indicates statistical significance ($\alpha$ = 0.05). (**C**) After 24 h, HCT116 cells were collected and resuspended in a fluorescent labeling mixture containing FITC-labeled Annexin-V. Cells were analyzed via flow cytometry. Cell counts collected from 3 biologically independent experiments (n = 3) were combined. Baseline cell death as a consequence of 24 h incubation in transfection reagent, as determined by analyzing cells transfected with RedTrackCMV (control vector) and incubated in the absence of *Cj*-CDT, was subtracted from the data represented by bars 3 and 4. Error bars represent standard deviations of data combined from 3 biologically independent experiments (n = 3). Statistical analyses of the data were conducted using one-way Anova followed by Dunnett's post hoc test. Data are presented as mean ± SD. $P < 0.05$ indicates statistical significance ($\alpha$ = 0.05).

### KEY RESOURCES TABLE

| REAGENT or RESOURCE | SOURCE | IDENTIFIER |
|---|---|---|
| **Antibodies** | | |
| Rabbit monoclonal α-LC3A/B (D3U4C) | Cell Signaling Technology, USA | Cat#12741; RRID:AB_2617131 |
| Mouse monoclonal α-β-Tubulin (D3U1W) | Cell Signaling Technology, USA | Cat#86298; RRID:AB_2715541 |
| Rabbit monoclonal α-γ-H2A.X (Ser139) (20E3) | Cell Signaling Technology, USA | Cat#9718; RRID:AB_2118009 |
| Rabbit monoclonal α-β-Actin (13E5) | Cell Signaling Technology, USA | Cat#4970; RRID:AB_2223172 |
| Rabbit monoclonal α-GAPDH (D16H11) | Cell Signaling Technology, USA | Cat#5174; RRID:AB_10622025 |
| Rabbit polyclonal α-α-Tubulin | Cell Signaling Technology, USA | Cat#2144; RRID:AB_2210548 |
| Mouse monoclonal α-Keima-Red | MBL International, Japan | Cat#M182-3M RRID:AB_10794910 |
| Rabbit polyclonal α-SQSTM1/p62 | Cell Signaling Technology, USA | Cat#5114; RRID:AB_10624872 |
| Rabbit polyclonal α-phospho-p53 (Ser15) | Cell Signaling Technology, USA | Cat#9284; RRID:AB_331464 |
| Mouse monoclonal α-p53 (DO-1) | Santa Cruz Biotechnology, USA | Cat#sc-126; RRID:AB_628082 |
| Rabbit monoclonal α-Ubiquitin (E4I2J) | Cell Signaling Technology, USA | Cat#43124; RRID:AB_2799235 |
| Mouse monoclonal α-Myc-Tag (9B11) | Cell Signaling Technology, USA | Cat#2276; RRID:AB_331783 |
| Rabbit monoclonal α-Myc-Tag (71D10) | Cell Signaling Technology, USA | Cat#2278; RRID:AB_490778 |
| Mouse monoclonal α-LC3B (E5Q2K) | Cell Signaling Technology, USA | Cat#83506; RRID:AB_2800018 |
| Rabbit monoclonal α-CtIP (D76F7) | Cell Signaling Technology, USA | Cat#9201; RRID:AB_10828593 |
| Rabbit polyclonal α-Cleaved Caspase-3 (Asp175) | Cell Signaling Technology, USA | Cat#9661; RRID:AB_2341188 |
| Rabbit monoclonal α-LC3A (D50G8) | Cell Signaling Technology, USA | Cat#4599; RRID:AB_10548192 |
| Rabbit monoclonal α-LC3B (D11) | Cell Signaling Technology, USA | Cat#3868; RRID:AB_2137707 |
| Rabbit monoclonal α-LC3C (D3O6P) | Cell Signaling Technology, USA | Cat#14736; RRID:AB_2798593 |
| Rabbit monoclonal α-GABARAP (E1J4E) | Cell Signaling Technology, USA | Cat#13733; RRID:AB_2798306 |
| Rabbit monoclonal α-GABARAPL1 (D5R9Y) | Cell Signaling Technology, USA | Cat#26632; RRID:AB_2798928 |
| Rabbit monoclonal α-GABARAPL2 (D1W9T) | Cell Signaling Technology, USA | Cat#14256; RRID:AB_2798436 |

| | | |
|---|---|---|
| Rabbit monoclonal α-p70 S6K (E8K6T) | Cell Signaling Technology, USA | Cat#34475 RRID:AB_2943679 |
| Rabbit polyclonal α-Phospho-p70 S6K (Thr389) | Cell Signaling Technology, USA | Cat#9205 RRID:AB_330944 |
| Rabbit monoclonal α-Phospho-ULK1 (Ser757) (D7O6U) | Cell Signaling Technology, USA | Cat#14202; RRID:AB_2665508 |
| Rabbit monoclonal α-ULK1 (D8H5) | Cell Signaling Technology, USA | Cat#:8054 RRID:AB_11178668 |
| Rabbit polyclonal α-Cj-CdtB | Yenzym Antibodies, USA | N/A |
| Goat α-Rabbit IgG (H+L), Biotinylated | Cell Signaling Technology, USA | Cat#14708; RRID:AB_2798581 |
| Goat polyclonal α-Rabbit IgG, HRP-linked | Cell Signaling Technology, USA | Cat#7074; RRID: AB_2099233 |
| Horse polyclonal α-Mouse IgG, HRP-linked | Cell Signaling Technology, USA | Cat#7076; RRID:AB_330924 |
| Goat α-biotin, HRP-linked | Cell Signaling Technology, USA | Cat#7075; RRID:AB_10696897 |
| Goat polyclonal α-Rabbit IgG (H+L), Alexa Fluor 488 Conjugated | Thermo Fisher Scientific, USA | Cat#A11008; RRID:AB_143165 |
| Donkey polyclonal α-Mouse IgG (H+L), Alexa Fluor 568 Conjugated | Thermo Fisher Scientific, USA | Cat#A10037; RRID:AB_2534013 |
| Protein A (HRP Conjugate) | Cell Signaling Technology, USA | Cat#12291 |
| **Bacterial and virus strains** | | |
| *Escherichia coli* BL21 (DE3) | New England Biolabs | Cat#C2527I |
| *Escherichia coli* DH5α | University of Illinois Urbana-Champaign Cell Media Facility | N/A |
| **Chemicals, peptides, and recombinant proteins** | | |
| 5-fluorouracil | Santa Cruz Biotechnology, USA | Cat#sc-29060; CAS: 51-21-8 |
| Actinomycin D | SelleckChem, Germany | Cat#S8964; CAS: 50-76-0 |
| Amicon Ultra-15 Centrifugal Filter Unit | Sigma-Aldrich, USA | Cat#UFC901024 |
| Bio-Gel P-60 Gel | Bio-Rad, USA | Cat#1504164 |
| Bleocin | Calbiochem, USA | Cat#203408; CAS: 55658-47-4 |
| Carbonyl-cyanide m-chlorophenyl hydrazine (CCCP) | Sigma-Aldrich, USA | Cat#C2759; CAS:555-60-2 |
| Cell Recovery Solution | Corning, USA | Cat#354253 |
| CHIR-99021 | Stem Cell Technologies, Canada | Cat#72052; CAS: 252917-06-9 |
| Chloroquine diphosphate (Kit component B) | Thermo Fisher Scientific, USA | Cat#P36240COMPONENTB; CAS: 50-63-5 |
| Click-iT L-homopropargylglycine (HPG) | Click Chemistry Tools, USA | Cat#1067 |

| | | |
|---|---|---|
| Cu-THPTA ligand | Lumiprobe, USA | Cat#H4050; CAS: 760952-88-3 |
| DAPI | Sigma-Aldrich, USA | Cat#D9542 |
| DEAE Sephacel | GE Healthcare, USA | Cat#17050001 |
| Dialysis Tubing 6000-8000 MWCO, Regenerated Cellulose | Fisher Scientific, USA | Cat#21-152-4 |
| Dimethyl sulfoxide | Sigma-Aldrich, USA | Cat# D2650; CAS: 67-68-5 |
| DMEM, high glucose, no glutamine, no methionine, no cystine | Thermo Fisher Scientific, USA | Cat#21013024 |
| DMEM/F-12 | Corning, USA | Cat#10-092-CV |
| Dulbecco's Modified Eagle Medium (DMEM) | Corning, USA | Cat#10-013-CV |
| E-64d | Sigma-Aldrich, USA | Cat#E8640; CAS: 88321-09-9 |
| Earl's Balanced Salt Solution (EBSS) | Thermo Fisher Scientific, USA | Cat#24010043 |
| Ethidium Homodimer-1 | Thermo Fisher Scientific, USA | Cat#L3224B; CAS: 61926-22-5 |
| Etoposide | Calbiochem, USA | Cat#341205; CAS: 33419-42-0 |
| FAM Picolyl Azide | Click Chemistry Tools, USA | Cat#1180 |
| Fetal Bovine Serum (FBS), Lot#16C315 | Sigma-Aldrich, USA | Cat#12306C |
| Gentle Cell Dissociation Reagent | Stem Cell Technologies, Canada | Cat#07174 |
| HALT Protease and Phosphatase Inhibitor Cocktail | Thermo Fisher Scientific, USA | Cat#78443 |
| HEPES | Gibco, USA | Cat#15630-080 |
| IntestiCult Mouse Organoid Growth Medium | Stem Cell Technologies, Canada | Cat#06005 |
| KU-55933 | Sigma-Aldrich, USA | Cat#118500; CAS: 587871-26-9 |
| Laemmli 6X SDS Loading Dye | Morganville Scientific, USA | Cat#LB0100 |
| Magnetic Dynabeads Protein A | Invitrogen | Cat#1001D |
| Matrigel | Corning, USA | Cat#354234 |
| McCoy's 5A (modified) | GE Healthcare, USA | Cat#SH30200.01 |
| MG132 | Sigma-Aldrich, USA | Cat#M7449 |
| N-ethylmaleimide | Sigma-Aldrich, USA | Cat#E1271; CAS: 128-53-0 |
| Nutlin-3 | Enzo Life Sciences, USA | Cat#ALX-430-128-M001; CAS: 548472-68-0 |
| Opti-MEM medium | Thermo Fisher Scientific, USA | Cat#11058021 |
| Penicillin-Streptomycin | Corning, USA | Cat#30-002-CI |
| Pepstatin A | Sigma-Aldrich, USA | Cat#P5318; CAS: 26305-03-3 |
| *Pfu*Ultra II Fusion HS DNA polymerase | Agilent Technologies, USA | Cat#600670 |
| Prolong Gold Antifade Reagent | Thermo Fisher Scientific, USA | Cat#P36930 |

| | | |
|---|---|---|
| Propidium Iodide | Thermo Fisher Scientific, USA | Cat#P3566 |
| Rapamycin | SelleckChem, Germany | Cat#S1039; CAS: 53123-88-9 |
| Recombinant *Campylobacter jejuni* cytolethal distending toxin | This Manuscript | N/A |
| Recombinant *Escherichia coli* cytolethal distending toxin | This Manuscript | N/A |
| Recombinant *Haemophilus ducreyi* cytolethal distending toxin | This Manuscript | N/A |
| RIPA Buffer | Thermo Fisher Scientific, USA | Cat#89901 |
| Rnase A | Sigma-Aldrich, USA | Cat#RNASEA-RO |
| Saponin-based permeabilization buffer (Component E of Click-iT Plus EdU Flow Cytometry Kit) | Thermo Fisher Scientific, USA | Cat#C10632 |
| Sodium L-ascorbate | Sigma-Aldrich, USA | Cat#11140; CAS: 134-03-2 |
| Staurosporine | Enzo Life Sciences, USA | Cat#ALX-380-014-C100; CAS: 62996-74-1 |
| SuperScript III Reverse Transcriptase | Thermo Fisher Scientific, USA | Cat#18080051 |
| SuperSignal West Femto Maximum Sensitivity Substrate | Thermo Fisher Scientific, USA | Cat#24095 |
| SuperSignal West Pico Plus Chemiluminescent Substrate | Thermo Fisher Scientific, USA | Cat#34580 |
| TAK-243 | SelleckChem, Germany | Cat#S8341; CAS: 1450833-55-2 |
| Talon Metal Affinity Resin (Cobalt) | Clontech Laboratories, USA | Cat#635503 |
| Universal Nuclease | Thermo Fisher Scientific, USA | Cat#88702 |
| Valproic Acid | Stem Cell Technologies. Canada | Cat#72292; CAS: 1069-66-5 |
| **Critical commercial assays** | | |
| MycoAlert Mycoplasma Detection Kit | Lonza, Switzerland | Cat#LT07-118 |
| BCA Protein Assay Kit | Thermo Fisher Scientific, USA | Cat#23225 |
| Lipofectamine 3000 kit | Thermo Fisher Scientific, USA | Cat#L300015 |
| Rneasy RNA Extraction Kit | Qiagen, Germany | Cat#74104 |
| Annexin-V PI Apoptosis Kit | BD Pharmingen, USA | Cat#556547 |
| **Experimental models: Cell lines** | | |
| Human: HCT116 | ATCC, USA | Cat#CCL-247 |
| Human: HCT116 ($p53^{+/+}$) | A gift from Dr. Bert Vogelstein | N/A |
| Human: HCT116 ($p53^{-/-}$) | A gift from Dr. Bert Vogelstein | RRID:CVCL_S744 |
| Human: HCT116 (RPS3-Keima) | A gift from Dr. Heeseon An | |

| | | |
|---|---|---|
| Human: HEK293T | ATCC, USA | Cat#CRL-3216 |
| Mouse: NIH/3T3 | A gift from Elizabeth Good | N/A |
| Mouse: Intestinal organoids (enteroids) from C57BL/6 | This Manuscript | N/A |
| Mouse: MEF (Atg5+/+) | A gift from Dr. Noboru Mizushima | |
| Mouse: MEF (Atg5-/-) | A gift from Dr. Noboru Mizushima | N/A |
| **Experimental models: Organisms/strains** | | |
| Mouse: C57BL/6 (for intestinal organoid harvest) | Breeding Colony, This Manuscript | N/A |
| **Oligonucleotides** | | |
| Primer: for cloning LC3B into RedTrackCMV; Forward: 5'-GAT ACT CGA GAT GCC GTC GGA GAA GAC CTT-3' | This Manuscript | N/A |
| Primer: for cloning LC3B into RedTrackCMV; Reverse: 5'-CTC AAG CTT TTA CAC TGA CAA TTT CAT CCC G-3' | This Manuscript | N/A |
| Primer: for point mutation of pET15b *Cj*-CdtB His-157 into Gly-157; Forward: 5'-GAT GCT TTT TTC AAT ATC GGT GCT TTA GCT AAT GG-3' | This Manuscript | N/A |
| Primer: for point mutation of pET15b *Cj*-CdtB His-157 into Gly-157; Reverse: 5'-CCA TTA GCT AAA GCA CCG ATA TTG AAA AAA GCA TC-3' | This Manuscript | N/A |
| Primer: *MAP1LC3B* qPCR; Forward: ACC ATG CCG TCG GAG AAG | Scherz-Shouval, *et al.*[1] | N/A |
| Primer: *MAP1LC3B* qPCR; Reverse: ATC GTT CTA TTA TCA CCG GGA TTT T | Scherz-Shouval, *et al.*[1] | N/A |
| Primer: *CDKN1A* qPCR; Forward: AGG CAC CGA GGC ACT CAG AG | Lynch and Milner[2] | N/A |
| Primer: *CDKN1A* qPCR; Reverse: AGT GGT AGA AAT CTG TCA TGC TG | Lynch and Milner[2] | N/A |
| Primer: *ACTB* qPCR; Forward: CAT GTA CGT TGC TAT CCA GGC | This Manuscript | N/A |
| Primer: *ACTB* qPCR; Reverse: CTC CTT AAT GTC ACG CAC GAT | This Manuscript | N/A |
| **Recombinant DNA** | | |
| Plasmid: pET-15b-CjCdtA | Eshraghi, *et al.*[3] | N/A |
| Plasmid: pET-15b-CjCdtB | Eshraghi, *et al.*[3] | N/A |
| Plasmid: pET-15b-CjCdtC | Eshraghi, *et al.*[3] | N/A |
| Plasmid: pET-15b-CjCdtB H157G | This Manuscript | N/A |
| Plasmid: pET-15b-EcCdtA | Eshraghi, *et al.*[3] | N/A |
| Plasmid: pET-15b-EcCdtB | Eshraghi, *et al.*[3] | N/A |
| Plasmid: pET-15b-EcCdtC | Eshraghi, *et al.*[3] | N/A |
| Plasmid: pET-15b-HdCdtA | Eshraghi, *et al.*[3] | N/A |
| Plasmid: pET-15b-HdCdtB | Eshraghi, *et al.*[3] | N/A |
| Plasmid: pET-15b-HdCdtC | Eshraghi, *et al.*[3] | N/A |
| Plasmid: pSelect-GFP-LC3B | InvivoGen, USA | Cat#psetz-gfplc3 |
| Plasmid: RedTrackCMV | RedTrackCMV was a gift from the lab of James Bamburg | Addgene Plasmid #50957; RRID:Addgene_50957 |

| | | |
|---|---|---|
| Plasmid: RedTrackCMV-LC3B | This Manuscript | N/A |
| Plasmid: pCMV-myc-LC3 | pCMV-myc-LC3 was a gift from the lab of Dr. Toren Finkel | Addgene Plasmid #24919; RRID:Addgene_24919 |
| **Software and algorithms** | | |
| SoftworX Explorer Suite, Version 3.5.1 | GE Life Sciences, USA | N/A |
| Imaris, Version 7.4.2 | Bitplane | N/A |
| Image Lab, Version 4.1 | Bio-Rad, USA | N/A |
| GraphPad Prism 7 | GraphPad, USA | N/A |
| FCS Express Version 6.06.0042 | De Novo Software, USA | N/A |
| Realplex, Version 2.2 | Eppendorf | N/A |
| **Other** | | |
| Lab-Tek II 8-well cell culture chamber slides | Thermo Fisher Scientific, USA | Cat#154534 |
| RT Widefield Microscope | GE Life Sciences, USA | N/A |
| CoolSnap HQ Camera | Photometrics | N/A |
| Vertical SDS-PAGE gel electrophoresis system | Bio-Rad, USA | Cat#1658004 |
| Wet Tank Western Blotting system | Bio-Rad, USA | Cat#1703930 |
| ChemiDoc XRS+ | Bio-Rad, USA | N/A |
| FACS Canto II Flow Cytometer | BD Biosciences, USA | N/A |
